# Niche-level immune evasion in *TP53* mutant AML residual disease revealed by spatial proteomics

**DOI:** 10.64898/2026.05.15.725421

**Authors:** Hideaki Mizuno, Yuki Nishida, Andrea Bedoy, Edward Ayoub, Yoonkyu Lee, Akshay Basi, Koji Sasaki, Guillermo Garcia-Manero, Jared Burks, Rashmi Kanagal-Shamanna, Michael Andreeff

## Abstract

Measurable (or minimal) residual disease (MRD) predicts relapse in patients with acute myeloid leukemia (AML). However, the biological and spatial characteristics of the AML bone marrow (BM) microenvironment (BMME) in which MRD cells survive remain largely unexplored; in particular, little is known of the BMME in *TP53* mutant (*TP53*^mut^) AML. Here, we applied sequential immunofluorescence to whole BM biopsy specimens obtained from patients with *TP53* wild-type (*TP53*^WT^) AML and *TP53*^mut^ AML at diagnosis and in morphological complete remission (CR) to generate a comprehensive spatial map of the hematopoietic and BMME components. We identified *TP53*^mut^ leukemia cells based on high p53 expression and delineated their spatial organization relative to stromal and immune niches. Biopsy-based cell composition analysis revealed marked B-cell depletion and an increased abundance of regulatory T-cells (Tregs) in *TP53*^mut^ BM at CR. Unlike *TP53*^WT^ BM, *TP53*^mut^ BM at CR exhibited persistent *TP53*^mut^ erythroid and immature leukemia cell clusters, spatially segregated from T-cell clusters, in perisinusoidal niches, suggesting niche-level immune evasion. Spatial profiling further revealed that Tregs characterized by FOXP3 upregulation were enriched near *TP53*^mut^ MRD cells, indicating a locally enhanced immunosuppressive activity. Single-cell RNA sequencing–based cell-cell communication analysis identified erythroid–T-cell interactions mediated by the GDF15-CD48 axis as a potential mechanism of T-cell suppression, suggesting that the erythroid differentiation of *TP53*^mut^ AML cells enhances local immunosuppression. Collectively, our results show a spatially organized immunosuppressive BMME in *TP53*^mut^ AML and highlight the potential of spatial proteomics to identify actionable MRD niches in leukemias.

**Key points:**

- *TP53* mutant erythroid and immature leukemia cells form spatial clusters segregated from T-cells in complete remission.
- An erythroblast-centered immunosuppressive niche characterizes *TP53* mutant leukemia cells.

## Introduction

Measurable (or minimal) residual disease (MRD) predicts relapse of hematological malignancies, including acute myeloid and lymphoblastic leukemias, regardless of detection by multicolor flow cytometry or mutation-oriented methods such as polymerase chain reaction and next generation-sequencing.^1–8^ Since relapsed leukemia cells are often resistant to conventional therapies and associated with short survival, understanding the biology of MRD is critical to developing first-line treatments that eradicate all disease in patients with acute myeloid leukemia (AML).^6,9^ Studies utilizing longitudinal single-cell RNA sequencing (scRNA-seq) of patient-derived specimens revealed that residual leukemia cells have upregulated quiescence and senescence signatures, potentially enabling them to survive therapy.^10,11^ However, the complex cell-cell interactions in the AML microenvironment that sustain MRD are largely unknown.

In AML patients, MRD is likely supported by the bone marrow (BM) microenvironment (BMME).^12^ Leukemia cells remodel the BMME and acquire treatment resistance through their interactions with non-hematopoietic BM cells, such as mesenchymal stromal cells (MSCs), through various pathways, including the SDF1/CXCL12, Wnt/β-catenin, VCAM/VLA-4/NF-κB, CD44, E-selectin, mitochondrial transfer, and TGFβ axes.^13–18^ An *in vivo* extramedullary BM system with genetically modified MSCs illustrated that HIF1α and NFκB are essential for AML engraftment.^19–21^ AML cells and MSCs also communicate with various immune cells to create an immune-permissive niche characterized by dysfunctional T and natural killer (NK) cells and an increase of immune-suppressive cells such as atypical B-cells, M2 macrophages, and myeloid-derived suppressor cells.^22,23^ This immunosuppressive microenvironment contributes to the limited efficacy of immunotherapies in AML compared to lymphoid malignancies.^22^ Advances in spatial proteomics and transcriptomics have facilitated the spatial mapping of complex structures in AML BM.^24,25^ For example, one study using spatial proteomics reported the proximity of NPM1c+ AML cells to MSCs or trabecular bone in post-treatment BM samples.^24^

*TP53* mutant (*TP53*^mut^) AML, which is characterized by poor treatment responses and high relapse rates and is associated with complex cytogenetic abnormalities,^26,27^ remains a major clinical challenge. Only 10-15% of patients with *TP53*^mut^ AML who are eligible for stem cell transplantation have long-term survival.^26,28,29^ Current MRD detection methods have limited predictive ability in *TP53*^mut^ AML compared to other AML subtypes, and *TP53*^mut^ MRD cells have not been well characterized.^7,26,30,31^ Immunohistochemistry-based MRD detection has high sensitivity and specificity, offering utility when BM aspiration is suboptimal, but its clinical predictive value has not been fully established and existing studies have not defined spatial architecture of *TP53*^mut^ AML.^32^

In the present study, subjecting BM biopsy specimens from healthy donors and AML patients to sequential immunofluorescence (seqIF) enabled the spatial mapping of malignant and non-malignant hematopoietic cells and the detailed characterization of *TP53*^mut^ MRD cells and their microenvironmental architecture. This analysis revealed dynamic alterations in the immune landscape, BM structures, and leukemia cells’ spatial organization and uncovered spatially segregated immune evasion mechanisms unique to *TP53*^mut^ AML. This study included the largest number of AML BM samples analyzed by spatial analyses thus far, defined targetable biological features of AML MRD, and demonstrated the broader potential of spatial MRD analysis across AML subtypes.

## Methods

### Patients and samples

Archival formalin-fixed, paraffin-embedded (FFPE) BM biopsy specimens were collected from the AML patients who were enrolled in clinical trials as part of MD Anderson’s AML “Moonshot” program and Break Through Cancer’s “Eradicating MRD in AML” project. Sample collection and analysis were performed under protocols approved by MD Anderson’s Institutional Review Board (PA19-0800/LAB02-395). Written informed consent was obtained from all patients in accordance with the Declaration of Helsinki.

### seqIF staining and imaging

FFPE tissues were cut into 5-µm sections and placed on slides. After preprocessing, the slides were loaded into the COMET system for seqIF.^33^ The output image files were subjected to data analysis.

### Data analysis

seqIF images were analyzed using Visiopharm (Visiopharm). BM regions of interest (ROI) underwent quality control to remove artifacts and low-quality areas. Nuclei were detected using a pretrained deep learning model, and cell boundaries were defined by nuclear expansion. Hematopoietic cell types were classified using percentile-based marker intensity thresholds optimized per marker and sample. Non-hematopoietic structural components were segmented based on marker expression and morphology. Protein expression was quantified using pixel intensities and normalized using Z-score^34^ or UniFORM-based approach.^35^ Spatial relationships were assessed using normalized median shortest distances (NMSD) between cell types, with permutation-based significance testing. Spatial neighborhood enrichment (SNE) was evaluated by clustering neighboring cells and testing enrichment using hypergeometric statistics.^24,36^ Structure proximity analysis for both cell types (SPA-cell) and clusters (SPA-cluster) was performed using a Poisson point process framework modeling cell-to-structure distances.^24^ scRNA-seq data were processed using Seurat,^37^ and cell-cell communication was analyzed using CellChat.^38^

More detailed methodologies are provided in the **supplemental Methods**.

## Results

### seqIF enables spatial mapping of whole BM specimens

Of the 139 *TP53*^WT^ and 105 *TP53*^mut^ AML patients we identified, 129 and 52, respectively, had CR. Sample selection was enriched for patients achieving CRs. Forty-seven randomly selected FFPE BM biopsy specimens (22 obtained at diagnosis and 25 at CR) from 10 *TP53*^WT^ and 19 *TP53*^mut^ AML patients, as well as 6 normal BM (NBM) biopsy specimens from healthy individuals, were subjected to seqIF. We selected NPM1 mutant BMs as TP53^WT^ to reduce heterogeneity of the TP53^WT^ group. Screening eliminated 24 specimens with insufficient staining and/or imaging quality (**supplemental Figure 1; supplemental Figure 2A-C**), yielding 16 *TP53*^mut^ AML BM samples (6 at diagnosis and 10 at CR), 10 *TP53*^WT^ AML BM samples (6 at diagnosis and 4 at CR), and 3 NBM samples for spatial proteomics analysis (**Figure 1A**). All AML patients whose samples were included in the analysis had received 1-4 courses of therapy at the time of CR sample collection (**Figure 1B; supplemental Table 1**).^39,40^ All of the 12 *TP53*^mut^ patients carried complex cytogenetic abnormalities, with 12/12 and 8/12 classified as biallelic *TP53* mutations by the International Consensus Classification and WHO criteria, respectively.^41,42^ Stringent tissue-quality filtering-based ROI selection mitigated the low signal-to-noise ratios resulting from the high autofluorescence of BM structures (**Figure 1C; supplemental Figure 3**; see supplemental Methods for details). Cell segmentation and supervised cell and structure classifications resulted in robust concordance between protein levels and cell and structure classification (**Figure 1D-G; supplemental Figure 4A-B**).

**Figure 1.**
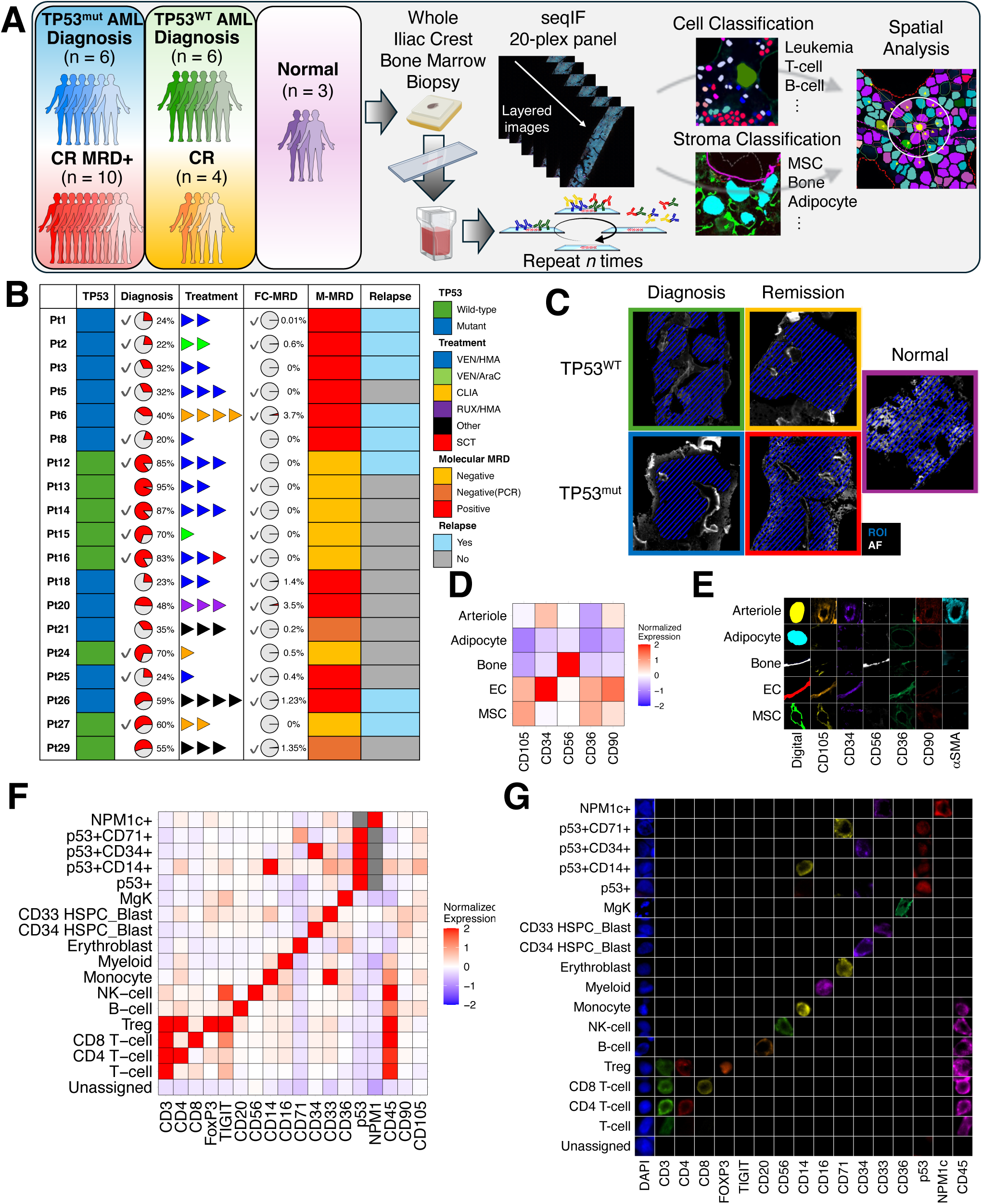
seqIF enables robust identification of hematopoietic cells and non-hematopoietic structures in BM biopsies. (A) Schematic workflow of the spatial proteomics analysis performed with the COMET system. (B) Patient characteristics. The pie charts in the “Diagnosis” and “FC-MRD” columns indicate the clinical smear blast count percentages and FC-MRD frequencies, respectively. The check marks in the columns indicate the 26 BM samples analyzed in the study. Treatment course details are shown in **supplemental Table 1**. M-MRD, molecular MRD; VEN, venetoclax; HMA, hypomethylating agent; AraC, cytarabine; CLIA, cladribine, idarubicin, and AraC; RUX, ruxolitinib; SCT, stem cell transplantation. (C) Representative seqIF images of ROI selection in BM samples. The blue shaded areas indicate the selected ROIs. AF, autofluorescence. (D) A heatmap showing the normalized protein expression of each structure in all BM samples (n = 29). The color scale indicates each structure’s mean marker Z-score. (E) Representative images showing each structure’s digital mask and marker expression. (F) A heatmap showing the normalized protein expression of each cell type in all analyzed samples (n = 29). The color scale indicates each cell type’s mean marker Z-score. (G) Representative seqIF images showing the marker expression of each cell type. Markers used for classification of each cell type are shown.

### seqIF and unbiased spatial analysis reveal spatial features of cells and structures in NBM

The cell type composition of NBM (**Figure 2A-B**) was consistent with the findings of previously reported spatial analyses of NBM.^24,43^ We calculated NMSDs to correct for differences in cellularity, with smaller values indicating greater proximity. Most cells were near cells of the same type; T-cell subsets were close to one another, and myeloid cells were spatially associated with monocytes (**Figure 2C; supplemental Figure 5A**). Unsupervised SNE analysis, a clustering method based on the similarity of each cell’s neighboring cell types (**supplemental Figure 6A**)^24,36^ showed that erythroblasts formed distinct clusters, consistent with erythroblastic island formation. Similarly, CD4 T-cells, CD8 T-cells, and B-cells (p=2.5×10^-39^, 5.6×10^-59^, and 4.8×10^-24^), monocytes and myelocytes (p=4.0×10^-20^ and <1×10^-300^), and hematopoietic stem progenitor cells (HSPCs; p=3.1×10^-8^) formed Ly, Mono/My/B/NK, and HSPC/T clusters, respectively, suggesting that these populations form functionally distinct niches within NBM (**Figure 2D**).

**Figure 2.**
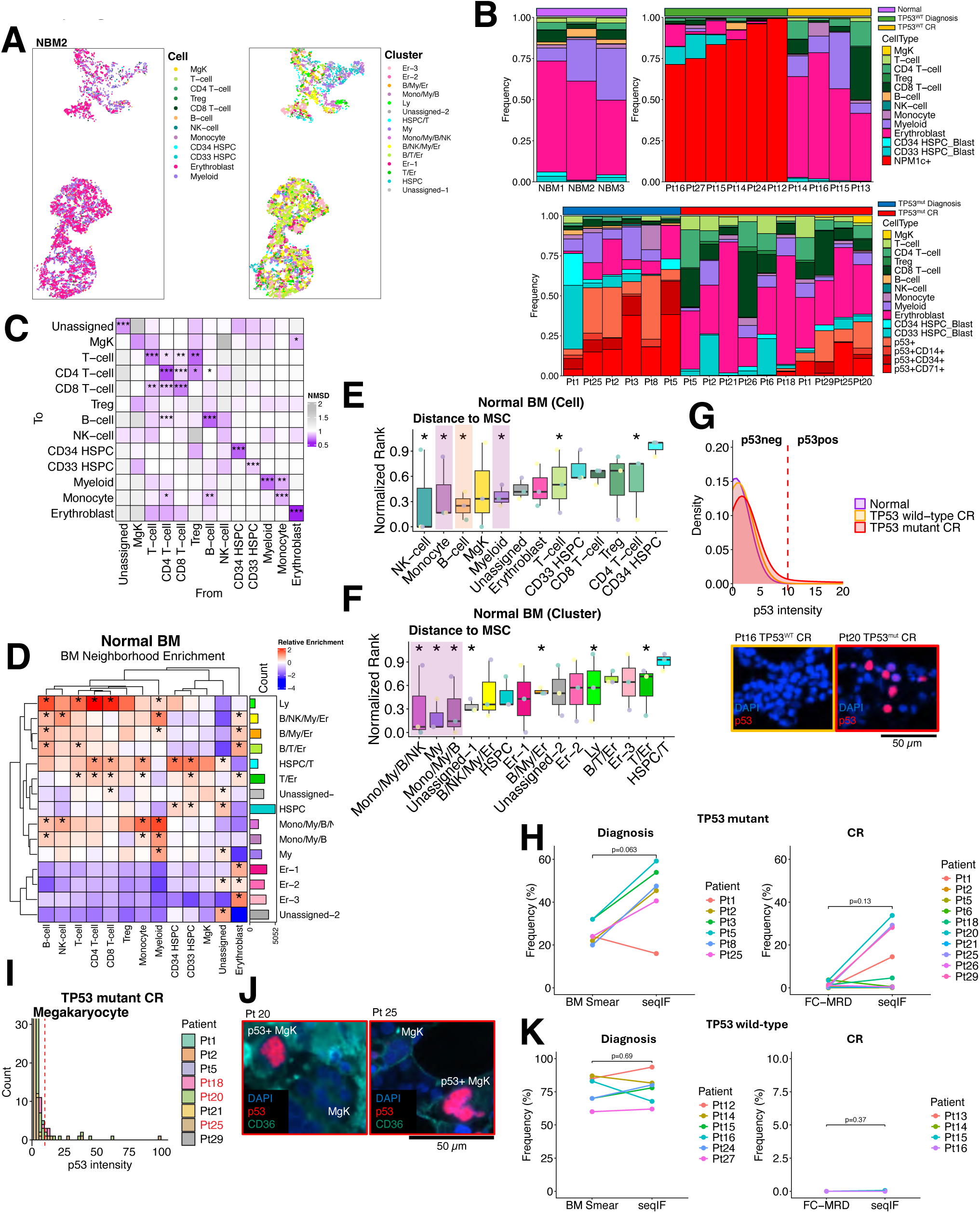
Spatial proteomics defines BM architecture and enables robust identification of *TP53*^WT^ and *TP53*^mut^ AML cells. (A) Dot plots showing cell types (left) and clusters (right) in a whole NBM sample from healthy donor 2 (NBM2). (B) Stacked bar plots showing the cell type compositions of all analyzed samples. The cell frequencies were calculated after removing unclassified cells. (C) A heatmap showing the NMSDs from the cell types in each column to the cell types in each row in NBM samples (n = 3). Empirical p-values were calculated by permuting the cell type labels and comparing the median distances in each image, and meta p-values were computed using the Stouffer method. *p < 0.05, **p < 0.01, ***p < 0.001. (D) A heatmap showing mean SNE in NBM samples (n = 3). The color scale indicates the enrichment of each cell type in each cluster relative to the entire population of that cell type. P-values were calculated using a hypergeometric test for the enrichment of each cell type, and adjusted p-values were computed using the Benjamini-Hochberg method. *Adjusted p < 0.05. (E, F) Boxplots of normalized rank distributions, showing the enrichment of each cell type (E) and each cluster (F) near MSC regions in NBM samples (n = 3). The dots indicate individual data points; horizontal lines, medians; boxes, interquartile ranges; whiskers extend to the most extreme values within 1.5 x interquartile ranges. Empirical p-values were calculated by permuting the cell type labels and comparing the median distances in each image, and meta p-values were calculated using the Stouffer method. *p < 0.05. (G) Top: A histogram showing p53 signal intensity in individual nuclei in NBM (n = 40,163 nuclei from n = 3 samples), *TP53*^WT^ BM at CR (n = 57,949 nuclei from n = 3 samples), and *TP53*^mut^ BM at CR (n = 112,142 nuclei from n = 10 samples). The red dashed vertical line at x = 10 indicates the threshold for p53 positivity. Bottom: Representative images of p53 staining in *TP53*^WT^ BM at CR from patient 16 (left) and *TP53*^mut^ BM at CR from patient 20 (right). (H) Dot plots showing leukemia cell frequencies in *TP53*^mut^ BM at diagnosis (left) and CR (right). Clinical smear-based blast percentages, FC-MRD frequencies, and leukemia cell frequencies measured by seqIF are shown. Data points for samples collected at the same times are connected by lines. P-values were calculated using the Wilcoxon rank-sum test. (I) A histogram showing the p53 signal intensities of megakaryocytes in *TP53*^mut^ BM at CR. The red dashed vertical line at x = 10 indicates the threshold for p53 positivity. The patients whose samples in which p53+ megakaryocytes were detected are labeled in red. (J) Representative images showing p53+ megakaryocytes (MgK) in *TP53*^mut^ BM at CR from patient 20 (left) and patient 25 (right). (K) Dot plots showing leukemia cell frequencies in *TP53*^WT^ BM at diagnosis (left) and CR (right). Clinical smear-based blast percentages, FC-MRD frequencies, and leukemia cell frequencies measured by seqIF are shown. Data points for samples collected at the same times are connected by lines. P-values were calculated using the Wilcoxon rank-sum test.

We next investigated spatial niches of individual cell types and structures. SPA-cell and SPA-cluster (**supplemental Figure 6B**) consistently showed spatial associations of B-cells, mature myeloid cells, and monocytes with MSCs (**Figure 2E-F; supplemental Figure 5B-C**). In contrast, HSPC clusters were near trabecular bone, with CD34 HSPC enrichment close to bone (p=2.4×10^-3^) (**supplemental Figure 7A-C**). This spatial arrangement aligns with previous findings that myeloid cells migrate from the endosteal to the sinusoidal niche during maturation.^24^ Erythroblasts were not preferentially associated with major structures, except that the Er-2 cluster was near endothelial cells (ECs), which is consistent with prior observations that erythroblastic islands are maintained by central macrophages rather than stromal structures^44^ (**Figure 2D-E; supplemental Figure 7A-B, D-G**). Collectively, these results demonstrate that seqIF combined with unbiased spatial analysis accurately recapitulates the architectural organization of NBM.

### seqIF enables the robust identification of *TP53*^mut^ and *TP53*^WT^ leukemia cells in BM

To detect *TP53*^mut^ leukemia cells, we leveraged the prior finding that AML cells harboring *TP53* missense mutations exhibit elevated p53 protein levels.^45,46^ Overall, cells in *TP53*^mut^ AML BM exhibited higher p53 levels than those in *TP53*^WT^ AML BM and NBM (p<2.2×10^-16^) (**supplemental Figure 8A**). With optimized thresholding, p53+ cells were detected in *TP53*^WT^ BM at CR with a false-positive rate lower than 0.01%, and no p53+ cells were detected in NBM (**Figure 2G; supplemental Table 4**). The two p53 false-positive cells (of 57,949 analyzed) in *TP53*^WT^ BM at CR were erythroblasts (**supplemental Figure 8B**).

p53+ leukemia cells were subclassified as p53+CD34 immature cells (p53-Im), p53+CD14+ monocyte-like cells (p53-Mo), p53+CD71+ erythroblast-like cells (p53-Er), and p53+CD34-CD14-CD71- unclassified cells (p53-U) (**supplemental Figure 4B**). Of the p53+ cells in *TP53*^mut^ BM at diagnosis and CR, 38.9% and 39.6%, respectively, were subclassified as p53-Er (**Figure 2B**). The frequencies of p53+ cells in *TP53*^mut^ BM biopsy samples tended to be higher than both clinical smear-based blast percentages at diagnosis (mean±SD, 43.8±15.1 vs. 25.7±5.13; p=0.063) and flow cytometry–based MRD (FC-MRD) frequencies at CR (5.33±6.48 vs. 1.24±1.35; p=0.13) (**Figure 2H**). Notably, seqIF detected MRD cells in patient 5’s *TP53*^mut^ BM at CR, which was negative for MRD by flow cytometry but positive for MRD by next-generation sequencing (**Figure 1B**). These discrepancies may reflect the absence of erythroid markers in routine FC-MRD panels.^1,4^ In 3 *TP53*^mut^ BM samples at CR, megakaryocytes exhibited high p53 expression, suggesting that they harbor *TP53* mutations (**Figure 2I,J**), which is aligned with a previous report linking *TP53* mutations with complex karyotypes to increased megakaryocyte progenitor abundance in *TP53*^mut^ AML.^47^ These results highlight the advantage of imaging-based approach, which can reliably identify megakaryocytes, a cell type often underrepresented in scRNA-seq datasets.^48^ Somatic *NPM1c* mutations in TP53^WT^ AML enabled leukemia cell identification with cytoplasmic localization and nuclear condensates of NPM1.^24,49,50^ The frequencies of leukemia cells detected by NPM1c closely matched the clinical smear-based blast percentages and FC-MRD frequencies (**Figure 2K**). seqIF detected NPM1c+ cells in two of four CR BM samples despite being negative for FC-MRD and molecular MRD (**supplemental Figure 8C**). Collectively, these results demonstrate that seqIF enables the robust identification of *TP53*^mut^ and *TP53*^WT^ *NPM1* mutant leukemia cells, facilitating the highly sensitive detection of MRD that standard clinical assays may miss.

### Biopsy-based cell composition analysis reveals a compromised immune microenvironment in *TP53*^mut^ AML BM at CR

Compared with *TP53*^WT^ BM at diagnosis, *TP53*^mut^ BM encompassed fewer leukemia cells (p=0.0022 (**Figure 3A**); a higher proportion of monocytes (p=0.015); and higher frequencies of mature myeloid cells (p=0.065) and erythroblasts (p=0.132) (**Figure 3B**). *TP53*^WT^ BM at CR was characterized by the expansion of multilineage cell components with an enrichment of erythroblasts, suggesting hematopoietic recovery, whereas *TP53*^mut^ BM lacked the reconstitution of hematopoietic lineages except for T-cells (**Figure 3B-C**). Compared to diagnostic samples, *TP53*^WT^ and *TP53*^mut^ BM at CR had fewer B-cells (p=0.026 and 0.067, respectively) (**Figure 3C**). In *TP53*^mut^ BM at CR, B-cells were frequently localized within sinusoids, suggesting their limited interactions with other immune cells within the marrow parenchyma (**Figure 3D**). Compared to *TP53*^WT^ BM, *TP53*^mut^ BM at both, diagnosis and CR had significant Treg enrichment (p=0.041 and p=0.031, respectively), indicating an immunosuppressive BMME (**Figure 3E**). In summary, these findings reveal distinct compositional and spatial immune alterations in *TP53*^mut^ BM, characterized by lower blast frequencies, myeloid enrichment, selective T-cell recovery, B-cell depletion, and increased Treg abundance, and collectively define a BMME with attenuated anti-leukemia immunity, a possible component of treatment resistance.

**Figure 3.**
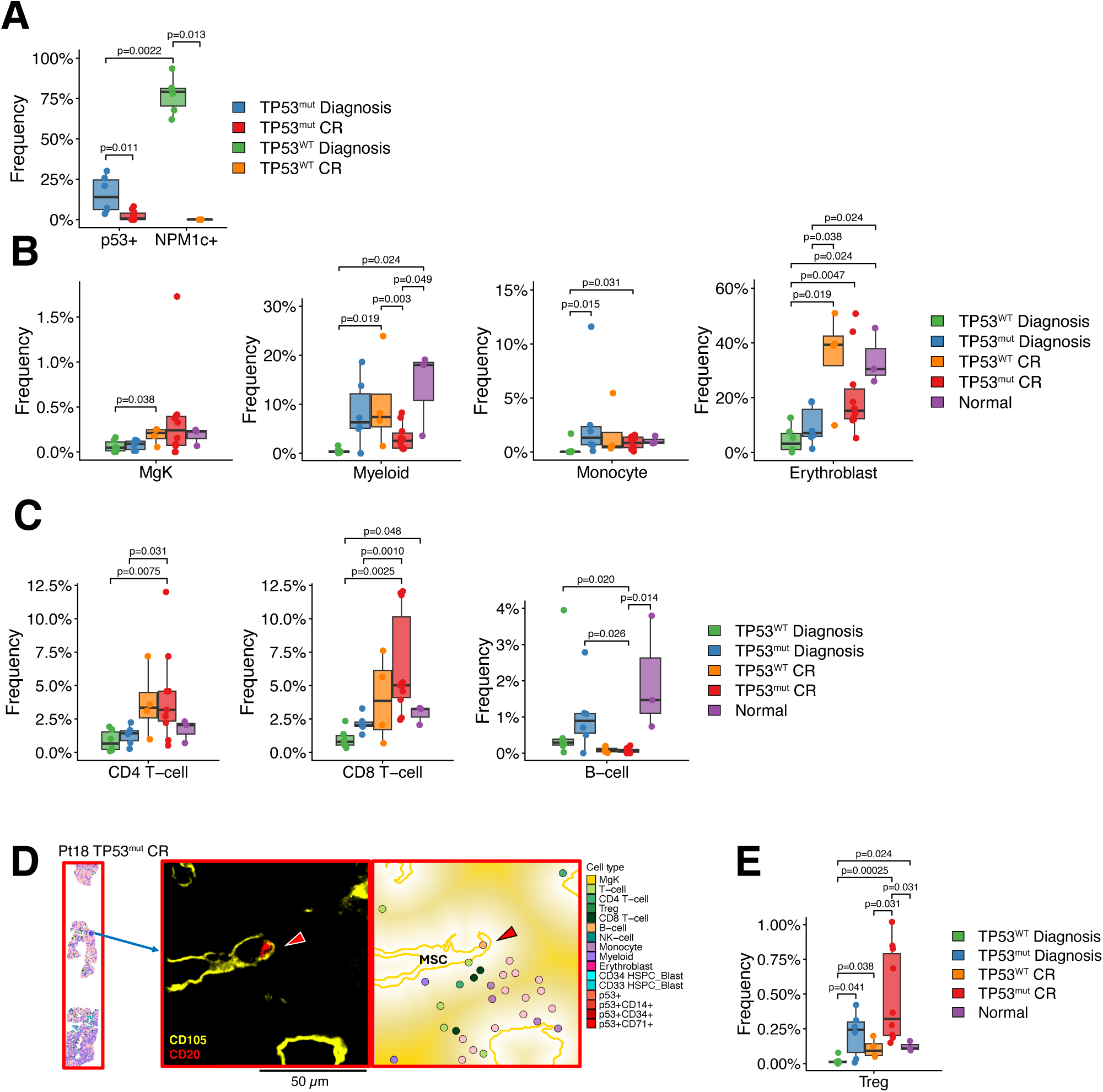
Biopsy-based BM composition analysis reveals a compromised immune microenvironment in *TP53*^mut^ AML. (A) Boxplot showing the frequencies of p53+ cells in *TP53*^mut^ BM samples at diagnosis (n = 6) and CR (n = 10) and frequencies of NPM1c+ cells in *TP53*^WT^ BM samples at diagnosis (n = 6) and CR (n = 4). The dots indicate individual data points; horizontal lines, medians; boxes, interquartile ranges; whiskers extend to the most extreme values within 1.5 x interquartile ranges. P-values were calculated using the Wilcoxon rank-sum test. Only significant comparisons (p<0.05) are shown. (B, C) Boxplots showing the frequencies of megakaryocytes (MgK), myeloid cells, monocytes, and erythroblasts (B) and frequencies of CD4 T-cells, CD8 T-cells, and B-cells (C) in *TP53*^WT^ BM samples at diagnosis (n = 6) and CR (n = 4), *TP53*^mut^ BM samples at diagnosis (n = 6) and CR (n = 10), and NBM samples (n = 3). The dots indicate individual data points; horizontal lines, medians; boxes, interquartile ranges; whiskers extend to the most extreme values within 1.5 x interquartile ranges. P-values were calculated using the Wilcoxon rank-sum test. Only significant comparisons (p<0.05) are shown. (D) Representative images showing B-cell colocalization in sinusoids. Left: A plot showing the cell types in whole *TP53*^mut^ BM at CR from patient 18. The blue rectangle indicates the area shown in the middle and right panels. Middle: A pseudo-color image of staining for CD105 (yellow) and CD20 (red). The red arrow indicates a B-cell. Right: A plot showing cells near MSC regions. The red arrow indicates a B-cell. The white-yellow background gradient indicates the cells’ distances from the MSC regions. (E) Boxplots showing the frequencies of Tregs in *TP53*^WT^ BM samples at diagnosis (n = 6) and CR (n = 4), *TP53*^mut^ BM samples at diagnosis (n = 6) and CR (n = 10), and NBM samples (n = 3). The dots indicate individual data points; horizontal lines, medians; boxes, interquartile ranges; whiskers extend to the most extreme values within 1.5 x interquartile ranges. P-values were calculated using the Wilcoxon rank-sum test. Only significant comparisons (p<0.05) are shown.

### *TP53*^mut^ AML BM has specialized clusters of multi-lineage p53+ cells that preferentially associate with MSCs

NMSD analysis revealed that, in both *TP53*^WT^ and *TP53*^mut^ BM at diagnosis, lymphocytes and erythroblasts preferentially localized with cells of the same lineage (**Figure 4A**). Similarly, SNE analysis identified distinct leukemia-, lymphocyte-, and erythroblast-specific clusters in *TP53*^WT^ and *TP53*^mut^ BM at diagnosis (**Figure 4B-C; supplemental Figure 9A**). NMSD analysis showed that p53-Im, p53-Mo, p53-Er, and p53-U formed their own clusters (p=1.6×10^-90^, 1.6×10^-67^, 5.7×10^-67^, and 1.6×10^-90^, respectively), resembling lineage-restricted hematopoietic populations (**Figure 4A; supplemental Figure 9B**). By SNE analysis, 38.2% of the *TP53*^WT^ BM clusters included the homogeneous NPM1c+ leukemia cluster (NPM1c), consisting of NPM1c+CD33+ leukemia cells, whereas the *TP53*^mut^ BM clusters were more heterogeneous, including p53CD34/CD71, p53CD34, p53CD14, p53CD71-1, p53CD71-2, and p53CD71-3 at frequencies ranging from 3.9% to 11.9% (**Figure 4C-D**). In *TP53*^mut^ BM, erythroblasts were intermingled with p53-Er within the p53CD71-1 and p53CD71-2 clusters (**Figure 4C**). This spatial overlap, also evident in the NMSD analysis (p=7.8×10^-47^) (**Figure 4A,E**), suggests that *TP53*^mut^ erythroid cells and normal erythroblasts occupy overlapping spatial niches. SPA-cell revealed that p53-U, p53-Er, and p53-Mo were predominantly near MSCs (p=5.0×10^-4^, 1.4×10^-4^, and 2.0×10^-41^, respectively) and, like HSPCs, p53-Im were closer to adipocytes (p=3.7×10^-5^) in *TP53*^mut^ BM at diagnosis (**Figure 4F; supplemental Figure 10; supplemental Figure 11A**). In contrast, NPM1c+ cells were not close to MSCs, suggesting interactions of *TP53*^mut^ but not NPM1c+ cells with MSCs (**Figure 4G**). SPA-cluster consistently showed p53CD34/CD71 clusters near MSCs (p=2.6×10^-14^), suggesting that p53-Im occupy two distinct stromal niches within *TP53*^mut^ BM at diagnosis (**Figure 4H-I; supplemental Figure 11B**). Similar to their localization in NBM, CD34+ HSPCs in *TP53*^mut^ BM were close to trabecular bone (p=1.4×10^-14^), suggesting that *TP53*^mut^ and *TP53*^WT^ CD34+ progenitor/stem cells rely on different specialized niches (**supplemental Figure 11C**). In both *TP53*^mut^ and *TP53*^WT^ BM at diagnosis, erythroblasts were near MSCs (p=7.9×10^-6^ and 1.65×10^-90^, respectively), suggesting an alternative stromal association of erythroblasts in the AML BMME (**Figure 4F-G**). Collectively, these findings demonstrate a *TP53*^mut^ AML–specific BMME that includes specialized clusters of multi-lineage p53+ cells and preferential associations of p53+ cells with MSCs.

**Figure 4.**
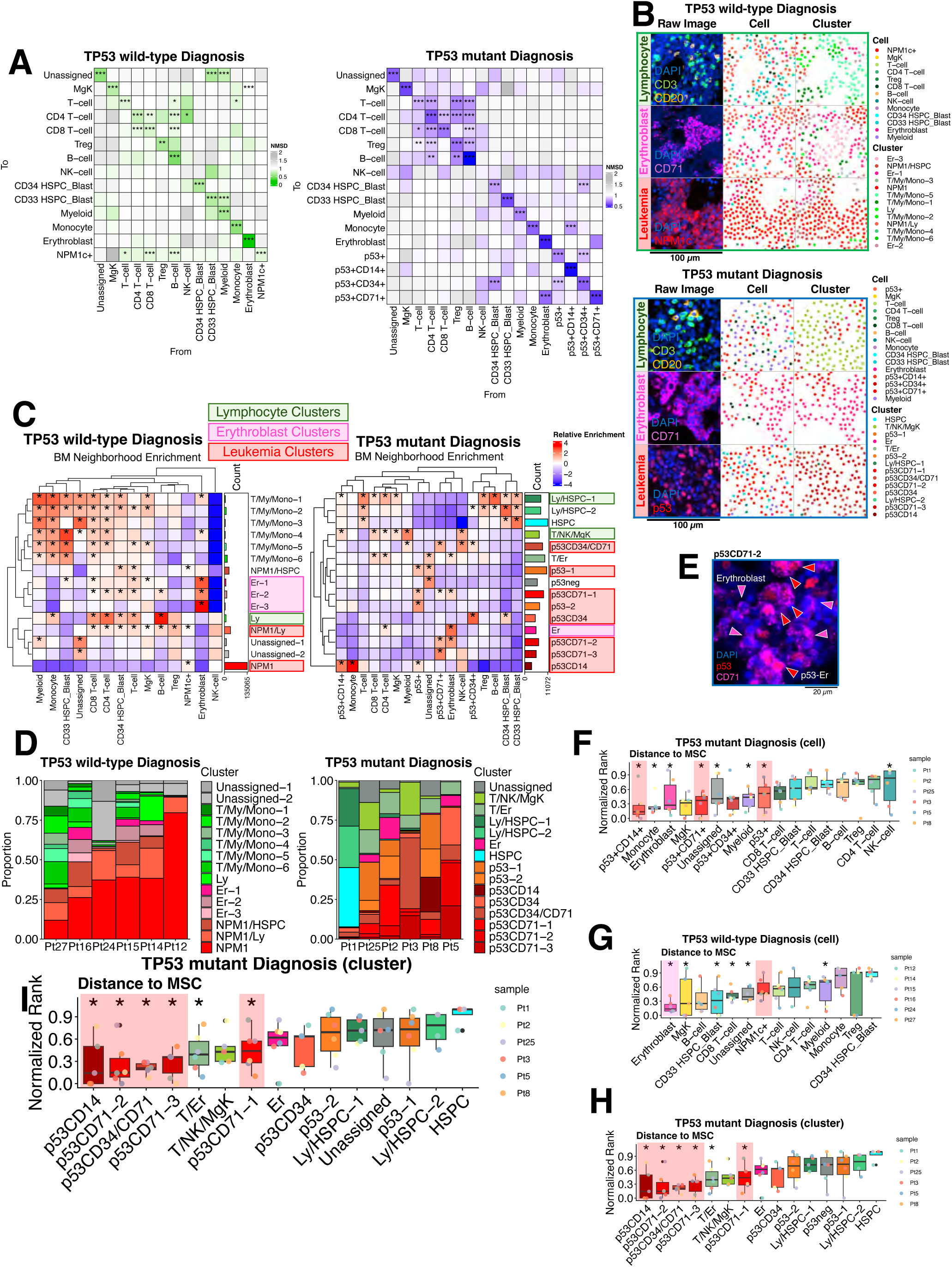
Spatial analysis of AML BM at diagnosis shows multi-lineage *TP53*^mut^ leukemia cell clusters in proximity to MSCs. (A) Heatmaps showing NMSDs from the cell types in each column to the cell types in each row in *TP53*^WT^ BM samples (n = 4; left) and *TP53*^mut^ BM samples (n = 6; right) at diagnosis. Empirical p-values were calculated by permuting the cell type labels and comparing the median distances within each image, and meta p-values were computed using the Stouffer method. *p < 0.05, **p < 0.01, ***p < 0.001. (B) Representative images of lymphocyte clusters, erythroblast clusters, and leukemia clusters in *TP53*^WT^ BM (top) and *TP53*^mut^ BM (bottom) at diagnosis. Pseudo-color images, cell type plots, and cluster plots are shown in the left, middle, and right columns, respectively. (C) Heatmaps showing SNE in *TP53*^WT^ BM samples (n = 6; left) and *TP53*^mut^ BM samples (n = 6; right) at diagnosis. The color scale indicates the enrichment of each cell type in each cluster relative to the entire population of that cell type. P-values were calculated using a hypergeometric test for the enrichment of each cell type, and adjusted p-values were calculated using the Benjamini-Hochberg method. *Adjusted p-value < 0.05. (D) Stacked bar plots showing the compositions of SNE clusters in *TP53*^WT^ BM (left) and *TP53*^mut^ BM (right) at diagnosis. (E) A representative seqIF image of a p53CD71-2 cluster. The red and pink arrows indicate p53+ erythroid cells and erythroblasts, respectively. (F-H) Boxplots of normalized rank distributions, showing the enrichment of each cell type near MSCs in *TP53*^mut^ BM samples (n = 6; F), each cell type near MSCs in *TP53*^WT^ BM samples (n = 6; G), and each cluster type near MSCs in *TP53*^mut^ BM samples (n = 6; H) at diagnosis. The dots indicate individual data points; horizontal lines, medians; boxes, interquartile ranges; whiskers extend to the most extreme values within 1.5 x interquartile ranges. Empirical p-values were computed by permuting the cell type labels and comparing the median distances in each image, and meta p-values were calculated using the Stouffer method. *p < 0.05. (I) Representative images showing p53CD34/CD71 clusters and MSC regions in *TP53*^mut^ BM at diagnosis. Left: A plot showing the cell types in whole *TP53*^mut^ BM at diagnosis from patient 5. The blue rectangle indicates the area shown in the right panels. Right: (i) a pseudo-color seqIF image of staining for DAPI (blue), p53 (red), CD34 (purple), and CD105 (yellow). (ii, iii) Plots showing cells labeled with cell type (ii) and cluster type (iii) near MSC regions. MSC regions are labeled with solid yellow lines. The white-yellow background gradient indicates the cells’ distances from the MSC regions.

### *TP53*^mut^ AML BM at CR is characterized by perisinusoidal niche-oriented *TP53*^mut^ erythroid-immature leukemia clusters segregated from lymphocyte clusters

Like in NBM and AML BM at diagnosis, lymphocytes, erythroblasts, and leukemia cells in AML BM at CR co-localized with their own cell type (**supplemental Figure 12A**). NMSD analysis revealed that p53-U, p53-Im, and p53-Er were in close proximity, with median distances of 121 µm between p53-U and p53-Im (p=6.8×10^-4^) and 58 µm between p53-Er and p53-U (p=5.4×10^-4^) (**Figure 5A**). *TP53*^mut^ BM at CR had two clusters consisting of p53-U, p53-Im, and p53-Er (p53/p53CD34/p53CD71-1 and -2) (**Figure 5B; supplemental Figure 12B**). A shorter NMSD between p53-Im and p53-Er in BM at CR compared to diagnosis (p=0.041) suggests a stronger spatial association of these cells at CR (**Figure 5C**). SPA-cluster revealed that the p53/p53CD34/p53CD71-1 cluster was close to MSCs (p=1.5×10^-22^), suggesting that the MSC niche supports *TP53*^mut^ MRD cells (**Figure 5D-E; supplemental Figure 13; supplemental Figure 14A**). *TP53*^WT^ BM at CR exhibited significant B-cell enrichment in the lymphocyte clusters Ly-1 (p=5.2×10^-12^) and Ly-2 (p=7.5×10^-6^), whereas *TP53*^mut^ BM did not, indicating reduced B–T-cell interaction in *TP53*^mut^ BM (**Figure 5B,F; supplemental Figure 14B-C**). Interestingly, whereas NPM1c+ cells clustered with CD4 and CD8 T-cells without Tregs, the T/HSPC/p53CD34 and T/HSPC/p53 clusters (which contained both p53+ cells and T-cells) included Tregs, suggesting a Treg-mediated immunosuppressive microenvironment surrounding residual p53+ cells (**Figure 5B**). SPA-cluster revealed lymphocyte clusters near sinusoids in *TP53*^WT^ BM (Ly-1; p=1.0×10^-34^) and *TP53*^mut^ BM (T-1; p=1.3×10^-24^) at CR, but not at diagnosis, suggesting the relocation of lymphocytes in BM after treatment (**Figure 4I**; **Figure 5D,F; supplemental Figure 11D; supplemental Figure 14B-C**). Although the p53/p53CD34/p53CD71-1 and T-1 clusters in *TP53*^mut^ BM at CR were in sinusoidal niches, cluster-wise NMSD analysis demonstrated that they were among the most spatially distant (p=4.6×10^-61^), highlighting the segregation of p53+ cells from lymphocyte clusters (**Figure 5G; supplemental Figure 14D-E**). In samples with both leukemia clusters (p53/p53CD34/p53CD71- 1 and -2) and lymphocyte clusters (T-1 and T-2), the clusters occupied distinct sinusoidal regions (p<0.005) (**Figure 5H-I**). Erythroblasts were less enriched near sinusoids, whereas mature myeloid cells and monocytes regained perisinusoidal localization in *TP53*^WT^ BM (p=8.6×10^-39^ and 1.9×10^-17^, respectively) and *TP53*^mut^ BM (p= 2.0×10^-77^ and 1.6×10^-27^) at CR, mirroring the architecture of NBM and suggesting anatomical recovery after treatment (**Figure 2E-F; supplemental Figure 14F-G**). In summary, *TP53*^mut^ BM at CR is characterized by perisinusoidal niche–oriented *TP53*^mut^ erythroid–immature leukemia clusters segregated from lymphocyte clusters, suggesting a *TP53*^mut^ AML–specific immunosuppressive BMME.

**Figure 5.**
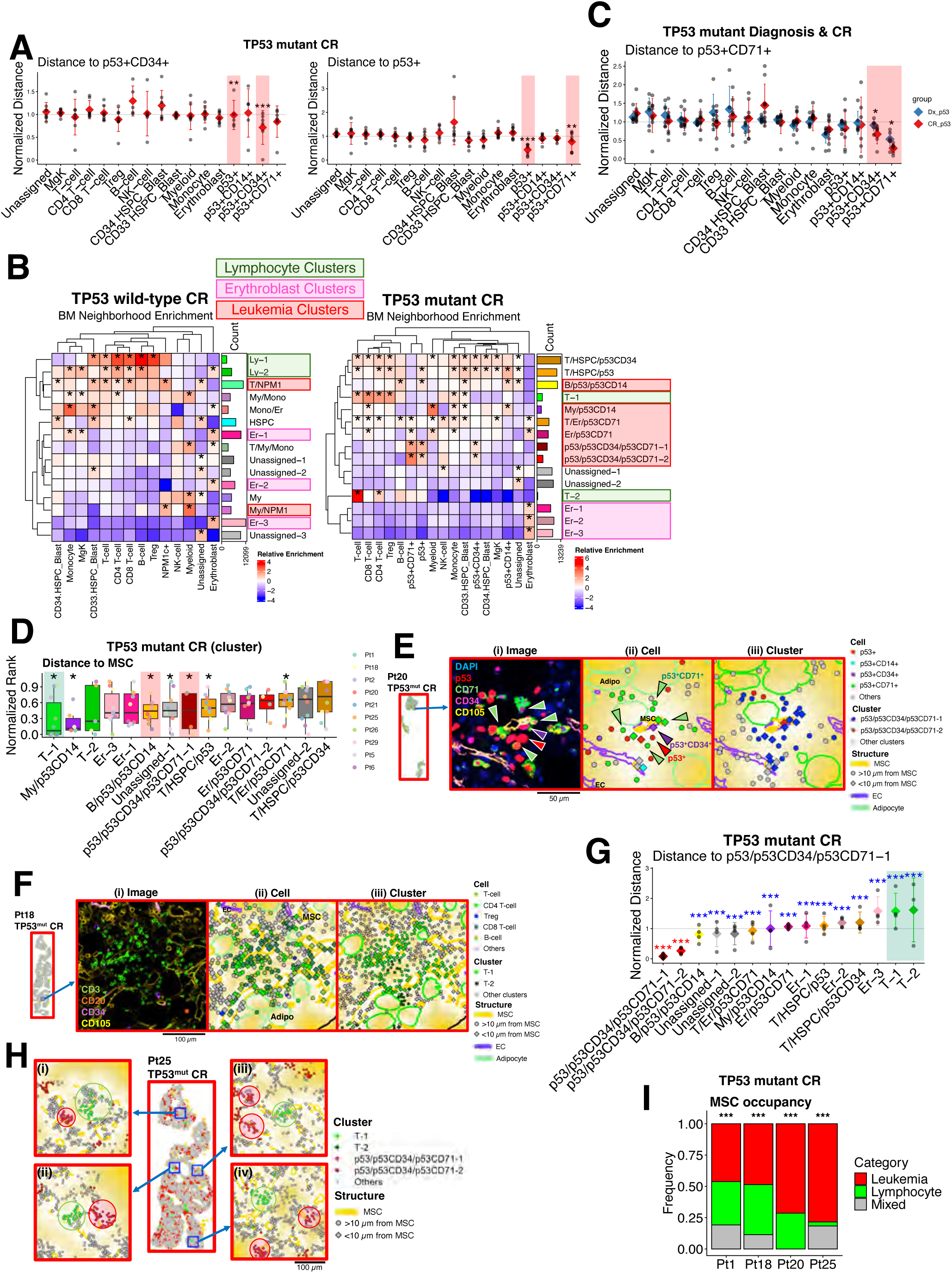
Spatial analysis of AML BM at CR reveals niche-level immune evasion in *TP53*^mut^ AML. (A) Plots showing different cell types’ NMSDs to p53+CD34+ leukemia cells (left) and p53+ leukemia cells (right) in *TP53*^mut^ BM samples at CR (n = 10). The dots indicate individual data points; diamonds, means; and error bars, standard deviations. Empirical p-values were calculated by permuting the cell type labels and comparing the median distances in each image, and meta p-values were calculated using the Stouffer method. *p < 0.05, **p < 0.01, ***p < 0.001. (B) Heatmaps of SNE in *TP53*^WT^ BM samples (n = 4; left) and *TP53*^mut^ BM samples (n = 10; right) at CR. The color scale indicates the enrichment of each cell type in each cluster relative to the entire population of that cell type. P-values were calculated using a hypergeometric test, and adjusted p-values were calculated using the Benjamini-Hochberg method. *Adjusted p-value < 0.05. Lymphocyte clusters, erythroblast clusters, and leukemia clusters are highlighted in green, pink, and red, respectively. (C) A plot showing different cell types’ NMSDs to p53+CD71+ leukemia cells in *TP53*^mut^ BM samples at diagnosis (n = 6; blue) and CR (n = 10; red). The dots indicate individual data points; diamonds, means; and error bars, standard deviations. P-values for differences between two groups in each cell type were calculated using the Wilcoxon rank-sum test and adjusted using the Benjamini-Hochberg method. *Adjusted p < 0.05. (D) Boxplots of normalized ranks of clusters, showing the enrichment of each cluster near MSCs in *TP53*^mut^ BM samples at CR (n = 10). The dots indicate individual data points; horizontal lines, medians; boxes, interquartile ranges; whiskers extend to the most extreme values within 1.5 x interquartile ranges. Empirical p-values were calculated by permuting the cell type labels and comparing the median distances in each image, and meta p-values were calculated using the Stouffer method. *p < 0.05. (E) Representative images of the p53/p53CD34/p53CD71-1 and -2 clusters in *TP53*^mut^ BM at CR from patient 20. Left: A plot showing the cell types in whole BM. The blue rectangle indicates the area shown in the right panels. Right: (i) a pseudo-color seqIF image of staining for DAPI (blue), p53 (red), CD71 (green), CD34 (purple), and CD105 (yellow). (ii, iii) Plots showing cells labeled with cell type (ii) and cluster type (iii) with structures. The white-yellow background gradient indicates the cells’ distances from the MSC regions. The red, purple, and green arrows indicate p53+ cells, p53+CD34+ cells, and p53+CD71+ cells, respectively. (F) Representative images of the T-1 and -2 clusters in *TP53*^mut^ BM at CR from patient 18. Left: A plot showing the cell types in whole BM. The blue rectangle indicates the area shown in the right panels. Right: (i) a pseudo-color seqIF image of staining for CD3 (green), CD20 (orange), CD34 (purple), and CD105 (yellow). (ii, iii) Plots showing cells labeled with cell type (ii) and cluster type (iii) with structures. The white-yellow background gradient indicates the cells’ distances from the MSC regions. (G) A plot showing different clusters’ NMSDs to cells in the p53/p53CD34/p53CD71-1 cluster in *TP53*^mut^ BM samples at CR (n = 10). The dots indicate individual data points; diamonds, means; and error bars, standard deviations. Empirical p-values were calculated by permuting the cell type labels and comparing the median distances in each sample, and meta p-values were calculated using the Stouffer method in each cluster. Three red asterisks (***) indicate a meta p < 0.001 for closeness to the p53/p53CD34/p53CD71-1 cluster. Three blue asterisks indicate a meta p < 0.001 for distantness to the p53/p53CD34/p53CD71-1 cluster. (H) Representative images of the segregation of the T-1 and p53/p53CD34/p53CD71-1 clusters in *TP53*^mut^ BM at CR from patient 25. Middle: A plot showing the cell types in whole BM. The blue rectangles indicate the area shown in the left and right panels. (i-iv) Plots showing cells labeled with cluster type with MSC regions. The white-yellow background gradient indicates the cells’ distances from the MSC regions. The red and green circles indicate T-1 clusters and p53/p53CD34/p53CD71-1 clusters adjacent to MSCs, respectively. (I) Bar plots showing the frequencies of MSC regions with leukemia clusters (p53/p53CD34/p53CD71-1 or p53/p53CD34/p53CD71-2), lymphocyte clusters (T-1 or T-2), or mixed enrichment in *TP53*^mut^ BM samples at CR from patients 1, 18, 20, and 25. P-values were calculated based on a permutation test. ***p < 0.001.

### *TP53*^mut^ MRD cells reside in an immunosuppressive, Treg-enriched niche in AML BM at CR

Across *TP53*^WT^ and *TP53*^mut^ BM samples whose leukemia cell frequencies were less than 1% (with mean frequencies of 0.062% and 0.29%, respectively), we identified 34 *TP53*^WT^ and 33 *TP53*^mut^ MRD cells (**Figure 6A**). The samples encompassed all four categories of p53+ cells, with p53-Er being the most frequent (**Figure 6B**), and 88.5% and 60.6% of the *TP53*^WT^ and *TP53*^mut^ MRD cells, respectively, were spatially isolated (p=0.012) (**Figure 6C**). The proportions of MRD cells accompanied by T-cells did not differ significantly between *TP53*^WT^ and *TP53*^mut^ BM (**Figure 6D**). However, *TP53*^mut^ MRD cells more frequently colocalized with Tregs (p=0.0060), suggesting microanatomical immunosuppression, whereas *TP53*^WT^ MRD cells were in a more inflammatory microenvironment (**Figure 6D-E; supplemental Figure 15A**). This difference became more pronounced when the analysis was restricted to isolated MRD cells (p=0.0026) (**supplemental Figure 15B**). Finally, both *TP53*^mut^ and *TP53*^WT^ MRD cells were adjacent to various structures, including MSCs, ECs, and adipocytes (**Figure 6F; supplemental Figure 15C-D)**. Collectively, these data suggest that after treatment, persistent *TP53*^mut^ MRD cells, unlike *TP53*^WT^ MRD cells, reside in an immunosuppressive, Treg-enriched niche.

**Figure 6.**
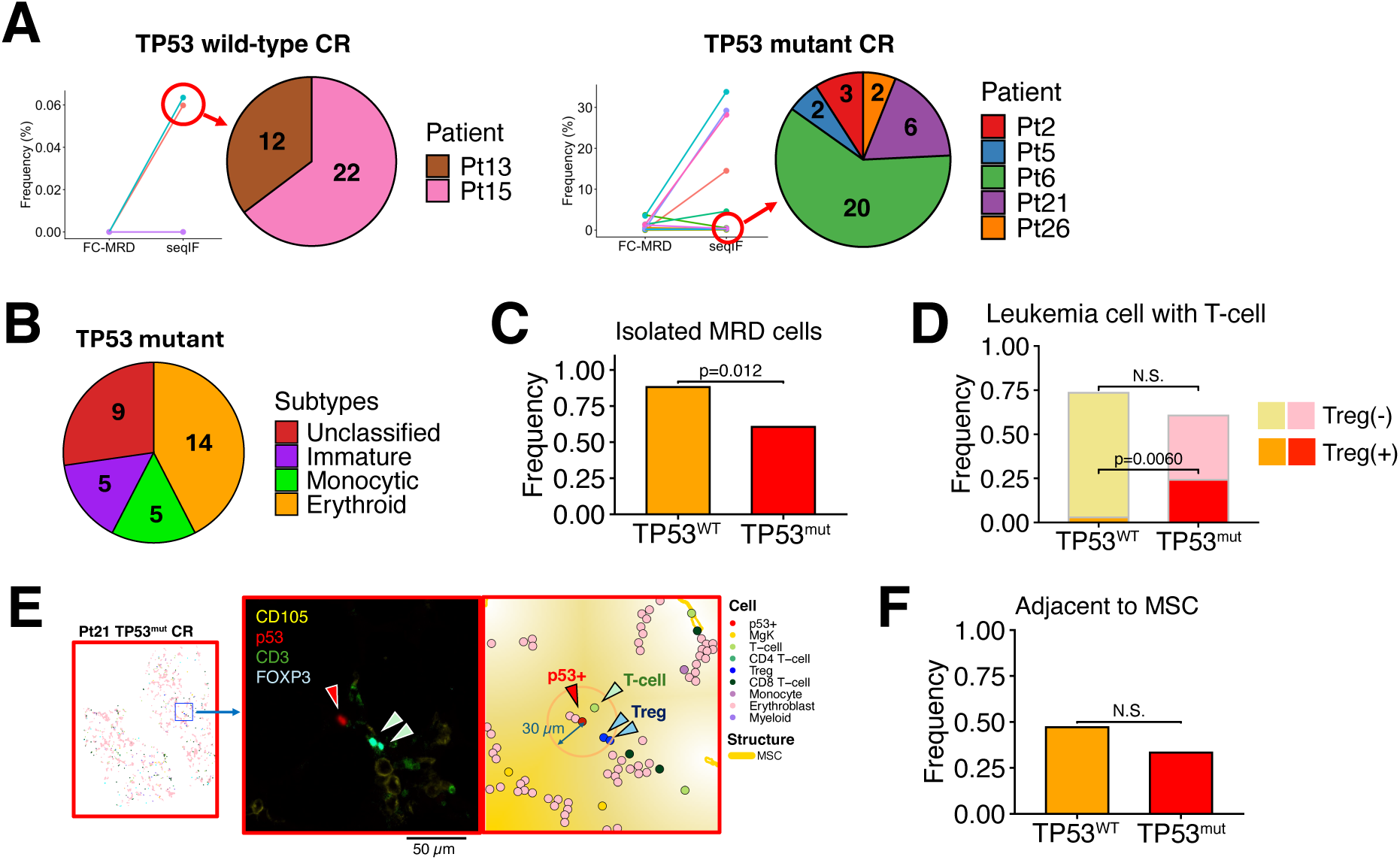
MRD-focused spatial analysis demonstrates colocalization of Treg with *TP53*^mut^ AML MRD cells. (A) Pie charts illustrating the patient sources of the *TP53*^WT^ (left) and *TP53*^mut^ (right) BM samples with leukemia cell frequencies of less than 1% in which seqIF detected MRD cells. Dot plots are adopted from Figure 2K (left) and 2H (right), showing frequencies of leukemia cells in BMs at CR. Red circles indicate samples included in pie charts. (B) A pie chart showing the p53+ subtypes of MRD cells in *TP53*^mut^ BM samples with leukemia cell frequencies of less than 1%. (C) Bar plots showing the frequencies of isolated MRD cells (defined as MRD cells without another leukemia cell within 30 μm) in *TP53*^WT^ BM samples (n = 34 MRD cells) and *TP53*^mut^ BM samples (n = 33 MRD cells). The p-value was calculated using the Fisher exact test. (D) Bar plots showing the frequencies of MRD cells associated with T-cells (defined as MRD cells within 30 µm of any T-cell) and MRD cells associated with Tregs (defined as MRD cells within 30 µm of any Treg) in *TP53*^WT^ BM samples (n = 34 MRD cells) and *TP53*^mut^ BM samples (n = 33 MRD cells). P-values were calculated using the Fisher exact test. N.S., not significant. (E) Representative images of a p53+ cell surrounded by T-cells in *TP53*^mut^ BM at CR from patient 21. Left: A plot showing the cell types in whole BM. The blue rectangle indicates the area shown in the middle and right panels. Middle: a pseudo-color seqIF image of staining for CD3 (green), p53 (red), FOXP3 (cyan), and CD105 (yellow). Right: Plots showing cells labeled with cell type with MSC regions. The white-yellow background gradient indicates the cells’ distances from the MSC regions. The green, blue, and red arrows indicate T-cells, Tregs, and p53+ cells, respectively. The pink circle in the right panel indicates the 30-µm distance from a p53+ cell. (F) Bar plots showing the frequencies of MRD cells adjacent to (within 10 µm of) MSCs in *TP53*^WT^ BM and *TP53*^mut^ BM. The p-value was calculated using the Fisher exact test.

### Distance-dependent dysfunctional T-cell immunity surrounds *TP53*^mut^ AML cells

To further investigate the immune microenvironment of *TP53*^mut^ BM, we examined spatially resolved phenotypic differences in T-cells, focusing on FOXP3 and TIGIT due to technical limitations precluding the detection of additional markers.^51,52^ High TIGIT levels are associated with suppressed effector function and reduced pro-inflammatory activity.^53–57^ Concordant with Treg-mediated suppression,^58^ CD4 T-cells and CD8 T-cells proximal to Tregs exhibited increased TIGIT levels than distant T-cells (p<2×10^-16^ and p=1.1×10^-15^, respectively) (**Figure 7A-C**), suggesting localized immunosuppressive interactions revealed by distance-based stratification.

**Figure 7.**
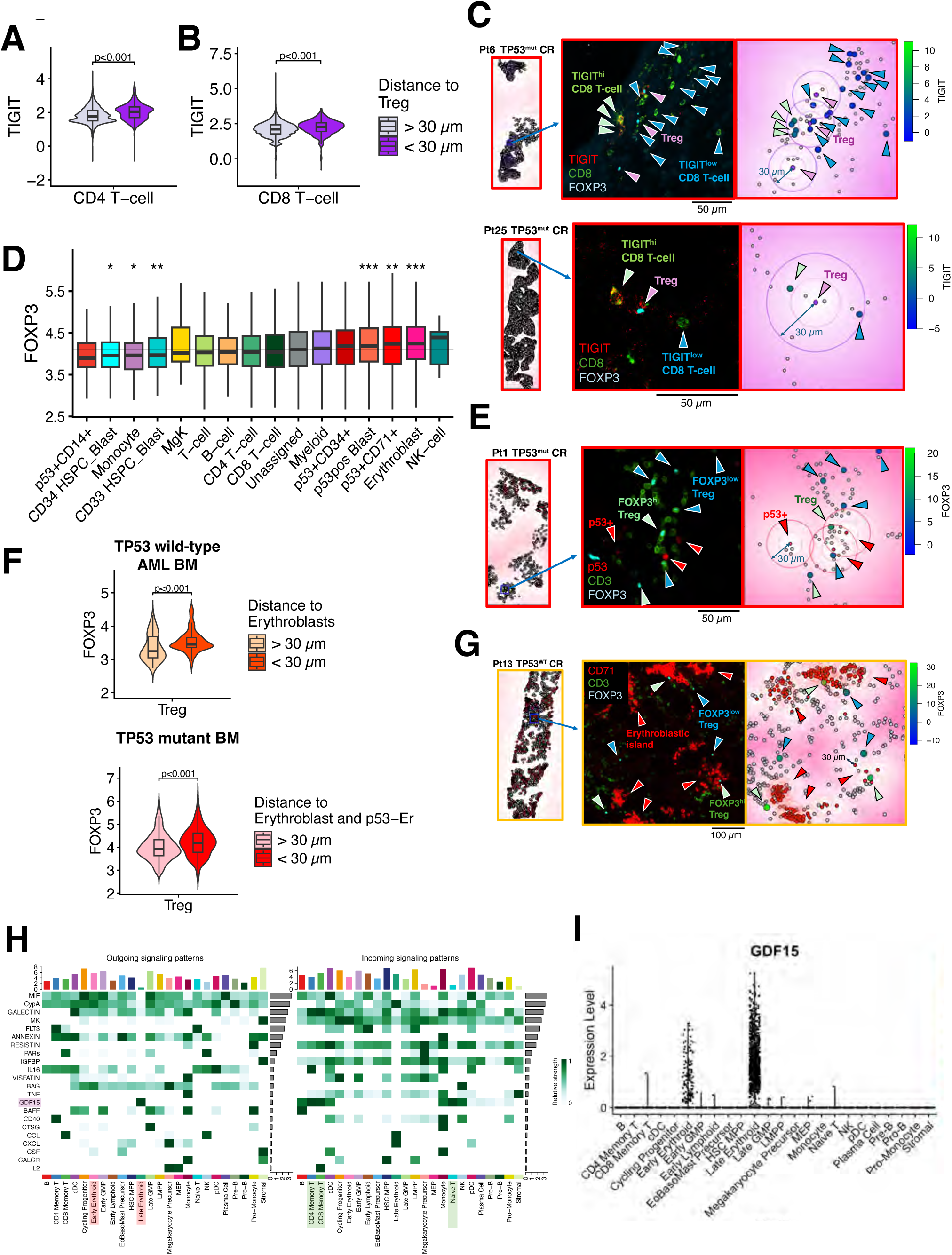
Distance-stratified protein expression analysis identifies a localized immunosuppressive microenvironment surrounding *TP53*^mut^ leukemia cells and erythroblasts. (A, B) Violin plots overlaid with boxplots showing the normalized TIGIT expression of CD4 T-cells <30 μm (n = 843) and >30 μm (n = 10,517) from Tregs (A) and CD8 T-cells <30 μm (n = 825) and >30 μm (n = 14,562) from Tregs (B) in all BM samples (n = 29). P-values were calculated using the Wilcoxon rank-sum test. (C) Representative images showing the TIGIT expression levels of CD8 T-cells near Tregs in *TP53*^mut^ BM at CR from patient 6 (top) and patient 25 (bottom). Left: plots showing the cell types in whole BMs. The blue rectangle indicates the area shown in the middle and right panels. Middle: a pseudo-color seqIF image of staining for TIGIT (red), CD8 (green), FOXP3 (cyan). Right: Plots showing cells. CD8 T-cells, Treg, and other cells are labeled with colors indicating normalized TIGIT expression, purple and grey, respectively. The color scales indicate each CD8 T-cell’s normalized TIGIT expression. The green, blue, and purple arrows indicate TIGIT^low^ CD8 T-cells, TIGIT^hi^ CD8 T-cells, and Tregs, respectively. The inner and outer circles indicate the 15-µm and 30-µm distances, respectively, from each Treg. In the dot plots, the white-purple background gradient indicates the distance from Tregs. (D) Boxplots showing the FOXP3 expression of Tregs close to each cell type. The horizontal lines indicate medians; boxes, interquartile ranges; and whiskers, 1.5 x interquartile ranges. The grey horizontal line spanning the graph indicates the median normalized FOXP3 expression of all Tregs (n = 615). P-values were calculated using a Wilcoxon rank-sum test comparing the normalized FOXP3 expression of Tregs <30 µm with that of Tregs >30 µm from each cell type, and adjusted p-values were calculated using the Benjamini-Hochberg method. *p < 0.05, **p < 0.01, ***p < 0.001. (E) Representative images showing FOXP3 levels in Tregs near any subtypes of p53+ cells in *TP53*^mut^ BM at CR from patient 1. Left: plots showing the cell types in whole BMs. The blue rectangle indicates the area shown in the middle and right panels. Middle: a pseudo-color seqIF image of staining for p53 (red), CD3 (green), FOXP3 (cyan). Right: Plots showing cells. Tregs, p53+ cells, and other cells are labeled with colors indicating normalized FOXP3 expression, red and grey, respectively. The color scale indicates each Treg’s normalized FOXP3 expression. The green, blue, and red arrows indicate FOXP3^low^ Tregs, FOXP3^hi^ Tregs, and p53+ cells, respectively. The inner and outer circles indicate the 15-µm and 30-µm distances, respectively, from each p53+ cell. In the dot plot, the white-red background gradient indicates the distance from the p53+ cells. (F) Violin plots overlaid with boxplots showing the normalized FOXP3 expression of Tregs <30 μm (n = 74) and >30 μm (n = 74) from erythroblasts in *TP53*^WT^ BM samples (n = 10; left) and Tregs <30 μm (n = 359) and >30 μm (n = 256) from erythroblasts in *TP53*^mut^ BM samples (n = 16; right). P-values were calculated using the Wilcoxon rank-sum test. (G) Representative images of FOXP3 levels in Tregs near erythroblasts in *TP53*^WT^ BM at CR from patient 13. Left: plots showing the cell types in whole BMs. The blue rectangle indicates the area shown in the middle and right panels. Middle: a pseudo-color seqIF image of staining for CD71 (red), CD3 (green), FOXP3 (cyan). Right: Plots showing cells. Tregs, erythroblasts, and other cells are labeled with colors indicating normalized FOXP3 expression, red and grey, respectively. The color scale indicates each Treg’s normalized FOXP3 expression. The green, blue, and red arrows indicate FOXP3^low^ Tregs, FOXP3^hi^ Tregs, and erythroblasts, respectively. The pink circles indicate the 30-μm distances from each erythroblast. In the dot plot, the white-red background gradient indicates the distance from the erythroblasts. (H) Heatmaps showing significantly enriched cell-cell communication pathways. Left: Outgoing signals based on the ligand expression. Right: Incoming signals based on the receptor expression. GDF15 in the row headings, erythroblasts in the column headings, and T-cells in the column headings are highlighted in purple, red, and green, respectively. (I) Violin plots of GDF15 expression in MD Anderson’s “Moonshot” scRNA-seq dataset.

Given that *TP53*^mut^ BMs are enriched with Tregs (**Figure 3C**), we hypothesized that Tregs near *TP53*^mut^ leukemia cells may have enhanced immunosuppressive properties. To test this, we compared the FOXP3 expression of Tregs close to each cell type with that of Tregs further from the same cell type across *TP53*^mut^ BM samples. Indeed, Tregs near p53-U and p53-Er exhibited higher FOXP3 levels compared to Tregs distant from these cells (p=5.6×10^-4^ and 3.4×10^-3^, respectively), suggesting that Tregs not only localize near leukemia cells but also actively display their immunosuppressive capacities (**Figure 7D-E**). Notably, Tregs near erythroblasts in both *TP53*^WT^ and *TP53*^mut^ BM had elevated FOXP3 levels (p=5.2×10^-4^ and 2.6×10^-6^, respectively) (**Figure 7D,F-G; supplemental Figure 16A-D**), consistent with recent reports identifying CD71+ erythroblasts as an emerging immunosuppressive cell.^59,60^ This suggests that *TP53*^mut^ leukemia cells hijack erythroid-mediated immunosuppression by differentiating into erythroid cells. In contrast, Tregs near erythroblasts in NBM did not exhibit high FOXP3 expression, suggesting an AML BM–specific mechanism (**supplemental Figure 16E**). Collectively, these findings suggest that Tregs in *TP53*^mut^ BM have enhanced immunosuppressive activity characterized by the distance-dependent upregulation of FOXP3, thereby contributing to an immunosuppressive BMME.

### The GDF15-CD48-Treg axis is a potential therapeutic target

To interrogate the mechanisms underlying the localized immunosuppression near erythroid cells, we analyzed our MD Anderson “Moonshot” scRNA-seq dataset, which includes 18 *TP53*^WT^ and 35 *TP53*^mut^ AML BM samples and 3 NBM samples.^61^ Cell-cell communication analysis revealed that erythroblasts communicate with T-cells through the GDF15-CD48 axis (p<0.001) (**Figure 7H; supplemental Table 6**). Prior studies identified a GDF15-CD48-Treg immunosuppression axis. The GDF15 concentrations in healthy donors (mean, 64.9 pg/ml), AML patients (mean, 550.4 pg/ml), and other cancer patients (median, 1902-3903 pg/ml), are too low to induce FOXP3 upregulation in Tregs.^62–65^ The observed upregulation suggests substantially high local concentration. GDF15 expression was almost exclusively restricted to erythroblasts (p<2.2×10^-308^ and p=1.5×10^-56^ for late and early erythroblasts, respectively), whereas CD48 was highly expressed in lymphocytes (p<2.2×10^-308^). These patterns were also observed in publicly available scRNA-seq datasets (**Figure 7I; supplemental Figure 17A-D**).^47,66^ Importantly, *TP53*^mut^, but not *TP53*^WT^, leukemia cells preferentially localize near erythroblasts and p53-Er (**Figure 4B**; **Figure 5B**), suggesting that this localized immunosuppressive niche selectively protects *TP53*^mut^ leukemia cells. Collectively, these findings indicate that erythroid cells create localized immunosuppressive niches in *TP53*^mut^ AML BM through GDF15-CD48-Treg signaling, providing mechanistic insight into the spatially restricted suppression of T-cell immunity.

## Discussion

Our study represents the largest spatial analysis of AML BM biopsy to date. Analyses of NBM recapitulated previously established spatial features, supporting the robustness of our approach.^24,43,44^ Spatial profiling of *TP53*^mut^ AML BM at diagnosis and at CR enabled characterization of MRD cells, providing insights beyond simple enumeration. Leveraging the NPM1c-specific staining pattern for *NPM1* mutations and high p53 protein levels for *TP53* missense mutations enabled us to confidently annotate *TP53*^WT^ and *TP53*^mut^ leukemia cells. Notably, *TP53*^mut^ AML BM showed enrichment of p53-Er, underscoring the importance of erythroid markers for identifying *TP53*^mut^ leukemia populations.^67^

Our findings indicate novel potential therapeutic targets in *TP53*^mut^ AML. First, we found that erythroid and immature leukemia cells formed perisinusoidal clusters that were spatially segregated from T-cell clusters, suggesting niche-level immune evasion, in *TP53*^mut^ BM at CR. MRD cells frequently colocalized with Tregs in *TP53*^mut^ BM, in contrast to the effector T-cell–dominant microenvironment in *TP53*^WT^ BM. Importantly, Tregs proximal to p53+ cells, particularly p53-Er, exhibited increased FOXP3 expression, indicating enhanced immunosuppressive activity. Prior single-cell analyses highlighted the enrichment of erythroid populations in *TP53*^mut^ AML^47,68^ but did not elucidate whether these cells contribute to therapy resistance. Our findings suggest that these *TP53*^mut^ erythroid cells actively establish localized immunosuppressive niches. Consistently, scRNA-seq cell-cell communication analysis identified GDF15-CD48-Treg as a potential mechanism of this immunosuppressive microenvironment. Although GDF15 is a well-established transcriptional target of p53,^69^ our results suggest that erythroid cells in AML BM commonly exploit additional regulatory mechanisms to upregulate GDF15. Thus, GDF15 represents a potential novel therapeutic target. Second, we observed increased Treg abundance in *TP53*^mut^ AML BM, which, given that TGFβ is a critical mediator of Treg induction, suggests that the TGFβ pathway is upregulated within the *TP53*^mut^ AML BMME. Indeed, we observed high phospho-SMAD3 expression in *TP53*^mut^ samples (**supplemental Figure 18**). The activation of the TGFβ pathway may also explain some key features of *TP53*^mut^ AML, including the enhanced erythroid differentiation and relatively lower leukemia burden. TGFβ signaling promotes the erythroid differentiation of HSPCs through the SMAD4-TIFγ axis,^70^ and *TP53* mutations may only partially impair TGFβ-mediated tumor suppressor functions, potentially resulting in a less proliferative phenotype.^71,72^ Thus, our findings indirectly support the notion that the *TP53*^mut^ BMME is characterized by TGFβ pathway upregulation, which may represent a therapeutically targetable vulnerability.

Our spatial proteomics analysis also uncovered previously unrecognized architectural and immunological features of the AML BMME. First, AML BM at CR had a striking depletion of B-cells. Such depletion may partially account for poor immunotherapy responses in AML patients, as intratumoral B-cell abundance is among the strongest predictors of immunotherapy response across solid tumors.^73–76^ Because BM aspirates are often contaminated by peripheral blood B-cells,^77^ biopsy-based assessment is essential to accurately evaluate B-cell depletion. Second, unlike NBM, in which erythroblastic islands are scaffolded by macrophages,^44^ erythroblasts in AML BM at diagnosis were preferentially localized to sinusoidal niches, suggesting that the leukemic microenvironment disrupts the macrophage-erythroblast interactions required for proper island formation.^44,78^ These findings reveal coordinated disruption of immune and erythroid niches in the AML BMME, potentially contributing to immune evasion and ineffective erythropoiesis.

Our use of a restricted antibody panel precluded the identification of certain immune populations, such as macrophages, plasma cells, and dendritic cells. In addition, leukemia cells harboring nonsense or frameshift *TP53* mutations, or *TP53* deletions resulting in the absence of p53 protein, cannot be detected by our method. Future studies incorporating expanded antibody panels and transcriptomic profiling will enable a comprehensive characterization of the AML BMME and facilitate therapeutic target discovery. In addition, our analysis was limited to two-dimensional sections of three-dimensional tissue, which may have led to the underrepresentation of z-stack spatial features. Validation using three-dimensional imaging approaches will be important to fully resolve BM organization.^79^

In conclusion, our study demonstrated that seqIF, by illustrating the spatial features of *TP53*^mut^ AML BM, such as the niche-level spatial dissociation of leukemia clusters from lymphocyte clusters and cell-level immune escape by accompanying activated Tregs, has utility in detecting and characterizing MRD cells. This framework is broadly applicable to the spatial analysis of solid tumors, offering a means to spatially dissect residual disease, tumor-immune interactions, and microenvironmental mechanisms underlying therapeutic resistance, thereby guiding the discovery and development of therapeutic targets.

## Acknowledgments

We thank the patients whose samples were used in the present study. This research was funded as part of the MD Anderson “Moonshot” program in AML and through Break Through Cancer’s “Eradicating MRD in AML” project (both to M.A.). Imaging and image analysis were performed in MD Anderson’s Flow Cytometry & Cellular Imaging Core Facility, supported by the National Institutes of Health (NIH) through MD Anderson’s Cancer Center Support Grant (2 P30 CA016672-48), a National Cancer Institute (NCI) Research Specialist Award (1 R50 CA243707-01A1; J.B.) and a Shared Instrumentation Award from the Cancer Prevention Research Institution of Texas (CPRIT). Slide preparation was performed by MD Anderson’s Research Histology Core Laboratory, which is supported in part by the NIH through MD Anderson’s Cancer Center Support Grant (2 P30 CA016672-48). H.M. was supported by a Takeda Science Foundation Overseas Research Fellowship. Additional funding was provided from the MD Anderson’s endowed Paul and Mary Haas Chair in Genetics (M.A.). E.A. and Y.L. are TRIUMPH Fellows in the CPRIT Training Program (RP210028).

## Authorship

Contribution: H.M., Y.N., and M.A. conceptualized and designed the study. H.M. collected imaging data in collaboration with A.B. and J.B. H.M. performed data analysis in collaboration with Y.N., E.A., Y.L., R.K.-S., J.B., and M.A. K.S., G.G.-M., and R.K.-S. provided specimens and pathology expertise. H.M., Y.N., and M.A. wrote the manuscript. All authors provided edits and approved the final manuscript.

## References

1. Heuser M, Freeman SD, Ossenkoppele GJ, et al. 2021 Update on MRD in acute myeloid leukemia: a consensus document from the European LeukemiaNet MRD Working Party. Blood. 2021;138(26):2753–2767.

2. Chandhok NS, Sekeres MA. Measurable residual disease in hematologic malignancies: a biomarker in search of a standard. EClinicalMedicine. 2025;86:103348.

3. Pantel K, Alix-Panabieres C. Minimal residual disease as a target for liquid biopsy in patients with solid tumours. Nat Rev Clin Oncol. 2025;22(1):65–77.

4. Cloos J, Valk PJM, Thiede C, et al. 2025 update on MRD in acute myeloid leukemia: a consensus document from the ELN-DAVID MRD Working Party. Blood. 2026;147(11):1147–1167.

5. Saygin C, Cannova J, Stock W, Muffly L. Measurable residual disease in acute lymphoblastic leukemia: methods and clinical context in adult patients. Haematologica. 2022;107(12):2783–2793.

6. Bazinet A, Kadia T, Short NJ, et al. Undetectable measurable residual disease is associated with improved outcomes in AML irrespective of treatment intensity. Blood Adv. 2023;7(13):3284–3296.

7. Morita K, Kantarjian HM, Wang F, et al. Clearance of Somatic Mutations at Remission and the Risk of Relapse in Acute Myeloid Leukemia. J Clin Oncol. 2018;36(18):1788–1797.

8. Sommer C, Mack HID, Killer MC, et al. Detection of minimal residual disease in circulating cell-free DNA in acute myeloid leukemia. Scientific Reports. 2025;15(1):32679.

9. Breems DA, Van Putten WL, Huijgens PC, et al. Prognostic index for adult patients with acute myeloid leukemia in first relapse. J Clin Oncol. 2005;23(9):1969–1978.

10. Li K, Du Y, Cai Y, et al. Single-cell analysis reveals the chemotherapy-induced cellular reprogramming and novel therapeutic targets in relapsed/refractory acute myeloid leukemia. Leukemia. 2023;37(2):308–325.

11. Duy C, Li M, Teater M, et al. Chemotherapy Induces Senescence-Like Resilient Cells Capable of Initiating AML Recurrence. Cancer Discov. 2021;11(6):1542–1561.

12. Kokkaliaris KD, Scadden DT. Cell interactions in the bone marrow microenvironment affecting myeloid malignancies. Blood Adv. 2020;4(15):3795–3803.

13. Jacamo R, Chen Y, Wang Z, et al. Reciprocal leukemia-stroma VCAM-1/VLA-4-dependent activation of NF-κB mediates chemoresistance. Blood. 2014;123(17):2691–2702.

14. Kim BR, Jung SH, Han AR, et al. CXCR4 Inhibition Enhances Efficacy of FLT3 Inhibitors in FLT3-Mutated AML Augmented by Suppressed TGF-b Signaling. Cancers (Basel). 2020;12(7):1737.

15. Zhou HS, Carter BZ, Andreeff M. Bone marrow niche-mediated survival of leukemia stem cells in acute myeloid leukemia: Yin and Yang. Cancer Biol Med. 2016;13(2):248–259.

16. Tabe Y, Shi YX, Zeng Z, et al. TGF-β-Neutralizing Antibody 1D11 Enhances Cytarabine-Induced Apoptosis in AML Cells in the Bone Marrow Microenvironment. PLoS One. 2013;8(6):e62785.

17. Jia Y, Zhang W, Basyal M, et al. FLT3 inhibitors upregulate CXCR4 and E-selectin ligands via ERK suppression in AML cells and CXCR4/E-selectin inhibition enhances anti-leukemia efficacy of FLT3-targeted therapy in AML. Leukemia. 2023;37(6):1379–1383.

18. Saito K, Zhang Q, Yang H, et al. Exogenous mitochondrial transfer and endogenous mitochondrial fission facilitate AML resistance to OxPhos inhibition. Blood Adv. 2021;5(20):4233–4255.

19. Jacamo R, Davis RE, Ling X, et al. Tumor Trp53 status and genotype affect the bone marrow microenvironment in acute myeloid leukemia. Oncotarget. 2017;8(48):83354–83369.

20. Chen Y, Jacamo R, Shi YX, et al. Human extramedullary bone marrow in mice: a novel in vivo model of genetically controlled hematopoietic microenvironment. Blood. 2012;119(21):4971–4980.

21. Battula VL, Le PM, Sun JC, et al. AML-induced osteogenic differentiation in mesenchymal stromal cells supports leukemia growth. JCI Insight. 2017;2(13):e90036.

22. Tettamanti S, Pievani A, Biondi A, Dotti G, Serafini M. Catch me if you can: how AML and its niche escape immunotherapy. Leukemia. 2022;36(1):13–22.

23. Lasry A, Nadorp B, Fornerod M, et al. An inflammatory state remodels the immune microenvironment and improves risk stratification in acute myeloid leukemia. Nat Cancer. 2023;4(1):27–42.

24. Bandyopadhyay S, Duffy MP, Ahn KJ, et al. Mapping the cellular biogeography of human bone marrow niches using single-cell transcriptomics and proteomic imaging. Cell. 2024;187(12):3120–3140.e3129.

25. Koedijk JB, van der Werf I, Penter L, et al. A multidimensional analysis reveals distinct immune phenotypes and the composition of immune aggregates in pediatric acute myeloid leukemia. Leukemia. 2024;38(11):2332–2343.

26. Grob T, Al Hinai ASA, Sanders MA, et al. Molecular characterization of mutant TP53 acute myeloid leukemia and high-risk myelodysplastic syndrome. Blood. 2022;139(15):2347–2354.

27. Badar T, Nanaa A, Atallah E, et al. Prognostic impact of ’multi-hit’ *versus* ’single-hit’ *TP53* alteration in patients with acute myeloid leukemia: results from the Consortium on Myeloid Malignancies and Neoplastic Diseases. Haematologica. 2024;109(11):3533–3542.

28. Shahzad M, Amin MK, Daver NG, et al. What have we learned about TP53-mutated acute myeloid leukemia? Blood Cancer J. 2024;14(1):202.

29. Stengel A, Kern W, Haferlach T, Meggendorfer M, Fasan A, Haferlach C. The impact of TP53 mutations and TP53 deletions on survival varies between AML, ALL, MDS and CLL: an analysis of 3307 cases. Leukemia. 2017;31(3):705–711.

30. Daver NG, Vyas P, Kambhampati S, et al. Tolerability and Efficacy of the Anticluster of Differentiation 47 Antibody Magrolimab Combined With Azacitidine in Patients With Previously Untreated AML: Phase Ib Results. J Clin Oncol. 2023;41(31):4893–4904.

31. Murdock HM, Kim HT, Denlinger N, et al. Impact of diagnostic genetics on remission MRD and transplantation outcomes in older patients with AML. Blood. 2022;139(24):3546–3557.

32. Brar N, Lawrence L, Fung E, et al. p53 immunohistochemistry as an ancillary tool for rapid assessment of residual disease in TP53-mutated acute myeloid leukemia and myelodysplastic syndromes. Am J Clin Pathol. 2024;162(3):269–281.

33. Rivest F, Eroglu D, Pelz B, et al. Fully automated sequential immunofluorescence (seqIF) for hyperplex spatial proteomics. Sci Rep. 2023;13(1):16994.

34. Hickey JW, Tan Y, Nolan GP, Goltsev Y. Strategies for Accurate Cell Type Identification in CODEX Multiplexed Imaging Data. Front Immunol. 2021;12:727626.

35. Wang K, Ait-Ahmad K, Kupp S, et al. Toward universal immunofluorescence normalization for multiplex tissue imaging with UniFORM. Cell Rep Methods. 2025;5(9):101172.

36. Schürch CM, Bhate SS, Barlow GL, et al. Coordinated Cellular Neighborhoods Orchestrate Antitumoral Immunity at the Colorectal Cancer Invasive Front. Cell. 2020;182(5):1341–1359.e1319.

37. Hao Y, Stuart T, Kowalski MH, et al. Dictionary learning for integrative, multimodal and scalable single-cell analysis. Nat Biotechnol. 2024;42(2):293–304.

38. Jin S, Plikus MV, Nie Q. CellChat for systematic analysis of cell-cell communication from single-cell transcriptomics. Nat Protoc. 2025;20(1):180–219.

39. Daver N, Senapati J, Kantarjian HM, et al. Azacitidine, Venetoclax, and Magrolimab in Newly Diagnosed and Relapsed Refractory Acute Myeloid Leukemia: Phase Ib/II Study and Correlative Analysis. Clin Cancer Res. 2025;31(12):2386–2398.

40. Kadia TM, Reville PK, Borthakur G, et al. Venetoclax plus intensive chemotherapy with cladribine, idarubicin, and cytarabine in patients with newly diagnosed acute myeloid leukaemia or high-risk myelodysplastic syndrome: a cohort from a single-centre, single-arm, phase 2 trial. Lancet Haematol. 2021;8(8):e552–e561.

41. Arber DA, Orazi A, Hasserjian RP, et al. International Consensus Classification of Myeloid Neoplasms and Acute Leukemias: integrating morphologic, clinical, and genomic data. Blood. 2022;140(11):1200–1228.

42. Hart SA, Lee LA, Seegmiller AC, Mason EF. Diagnosis of TP53-mutated myeloid disease by the ICC and WHO fifth edition classifications. Blood Adv. 2025;9(3):445–454.

43. Sarachakov A, Varlamova A, Svekolkin V, et al. Spatial mapping of human hematopoiesis at single-cell resolution reveals aging-associated topographic remodeling. Blood. 2023;142(26):2282–2295.

44. Chasis JA, Mohandas N. Erythroblastic islands: niches for erythropoiesis. Blood. 2008;112(3):470–478.

45. Tashakori M, Kadia T, Loghavi S, et al. TP53 copy number and protein expression inform mutation status across risk categories in acute myeloid leukemia. Blood. 2022;140(1):58–72.

46. Saft L, Karimi M, Ghaderi M, et al. p53 protein expression independently predicts outcome in patients with lower-risk myelodysplastic syndromes with del(5q). Haematologica. 2014;99(6):1041–1049.

47. Zeng AGX, Iacobucci I, Shah S, et al. Single-cell Transcriptional Atlas of Human Hematopoiesis Reveals Genetic and Hierarchy-Based Determinants of Aberrant AML Differentiation. Blood Cancer Discov. 2025;6(4):307–324.

48. Sun S, Jin C, Si J, et al. Single-cell analysis of ploidy and the transcriptome reveals functional and spatial divergency in murine megakaryopoiesis. Blood. 2021;138(14):1211–1224.

49. Falini B, Nicoletti I, Martelli MF, Mecucci C. Acute myeloid leukemia carrying cytoplasmic/mutated nucleophosmin (NPMc+ AML): biologic and clinical features. Blood. 2007;109(3):874–885.

50. Datar GK, Khabusheva E, Anand A, et al. Disparate leukemia mutations converge on nuclear phase-separated condensates. Cell. 2025;188(25):7118–7136.e7121.

51. Waldman AD, Fritz JM, Lenardo MJ. A guide to cancer immunotherapy: from T cell basic science to clinical practice. Nat Rev Immunol. 2020;20(11):651–668.

52. Sallman DA, McLemore AF, Aldrich AL, et al. TP53 mutations in myelodysplastic syndromes and secondary AML confer an immunosuppressive phenotype. Blood. 2020;136(24):2812–2823.

53. Chauvin JM, Zarour HM. TIGIT in cancer immunotherapy. J Immunother Cancer. 2020;8(2):e000957.

54. Johnston RJ, Comps-Agrar L, Hackney J, et al. The immunoreceptor TIGIT regulates antitumor and antiviral CD8(+) T cell effector function. Cancer Cell. 2014;26(6):923–937.

55. Joller N, Lozano E, Burkett PR, et al. Treg cells expressing the coinhibitory molecule TIGIT selectively inhibit proinflammatory Th1 and Th17 cell responses. Immunity. 2014;40(4):569–581.

56. Lai Y, Wang S, Ren T, et al. TIGIT deficiency promotes autoreactive CD4(+) T-cell responses through a metabolicLepigenetic mechanism in autoimmune myositis. Nat Commun. 2025;16(1):4502.

57. Zhang Q, Bi J, Zheng X, et al. Blockade of the checkpoint receptor TIGIT prevents NK cell exhaustion and elicits potent anti-tumor immunity. Nat Immunol. 2018;19(7):723–732.

58. Imianowski CJ, Chen Q, Workman CJ, Vignali DAA. Regulatory T cells in the tumour microenvironment. Nat Rev Cancer. 2025;25(9):703–722.

59. Elahi S, Ertelt JM, Kinder JM, et al. Immunosuppressive CD71+ erythroid cells compromise neonatal host defence against infection. Nature. 2013;504(7478):158–162.

60. Elahi S, Mashhouri S. Immunological consequences of extramedullary erythropoiesis: immunoregulatory functions of CD71(+) erythroid cells. Haematologica. 2020;105(6):1478–1483.

61. Li L, Muftuoglu M, Ayoub E, et al. Somatic TP53 Mutations Drive T and NK Cell Dysfunction in AML and Can be Rescued by Reactivating Wild Type p53. medRxiv. 2025:2025.2004.2011.25325281.

62. Hegab HM, El-Ghammaz AMS, El-Razzaz MK, Helal RAA. Prognostic Impact of Serum Growth Differentiation Factor 15 Level in Acute Myeloid Leukemia Patients. Indian J Hematol Blood Transfus. 2021;37(1):37–44.

63. Al-Sawaf O, Weiss J, Skrzypski M, et al. Body composition and lung cancer-associated cachexia in TRACERx. Nat Med. 2023;29(4):846–858.

64. Groarke JD, Crawford J, Collins SM, et al. Ponsegromab for the Treatment of Cancer Cachexia. N Engl J Med. 2024;391(24):2291–2303.

65. Wang Z, He L, Li W, et al. GDF15 induces immunosuppression via CD48 on regulatory T cells in hepatocellular carcinoma. J Immunother Cancer. 2021;9(9):e002787.

66. van Galen P, Hovestadt V, Wadsworth Ii MH, et al. Single-Cell RNA-Seq Reveals AML Hierarchies Relevant to Disease Progression and Immunity. Cell. 2019;176(6):1265–1281 e1224.

67. Robinson TM, Bowman RL, Persaud S, et al. Single-cell genotypic and phenotypic analysis of measurable residual disease in acute myeloid leukemia. Sci Adv. 2023;9(38):eadg0488.

68. Rodriguez-Meira A, Norfo R, Wen S, et al. Single-cell multi-omics identifies chronic inflammation as a driver of TP53-mutant leukemic evolution. Nat Genet. 2023;55(9):1531–1541.

69. Osada M, Park HL, Park MJ, et al. A p53-type response element in the GDF15 promoter confers high specificity for p53 activation. Biochem Biophys Res Commun. 2007;354(4):913–918.

70. Blank U, Karlsson S. TGF-β signaling in the control of hematopoietic stem cells. Blood. 2015;125(23):3542–3550.

71. Cordenonsi M, Dupont S, Maretto S, Insinga A, Imbriano C, Piccolo S. Links between tumor suppressors: p53 is required for TGF-beta gene responses by cooperating with Smads. Cell. 2003;113(3):301–314.

72. Kadia TM, Jain P, Ravandi F, et al. TP53 mutations in newly diagnosed acute myeloid leukemia: Clinicomolecular characteristics, response to therapy, and outcomes. Cancer. 2016;122(22):3484–3491.

73. Chang TG, Spathis A, Schäffer AA, et al. Tumor and blood B-cell abundance outperforms established immune checkpoint blockade response prediction signatures in head and neck cancer. Ann Oncol. 2025;36(3):309–320.

74. Griss J, Bauer W, Wagner C, et al. B cells sustain inflammation and predict response to immune checkpoint blockade in human melanoma. Nat Commun. 2019;10(1):4186.

75. Helmink BA, Reddy SM, Gao J, et al. B cells and tertiary lymphoid structures promote immunotherapy response. Nature. 2020;577(7791):549–555.

76. Restelli C, Ruella M, Paruzzo L, Tarella C, Pelicci PG, Colombo E. Recent Advances in Immune-Based Therapies for Acute Myeloid Leukemia. Blood Cancer Discov. 2024;5(4):234–248.

77. Waidhauser J, Schuh A, Trepel M, Schmälter AK, Rank A. Chemotherapy markedly reduces B cells but not T cells and NK cells in patients with cancer. Cancer Immunol Immunother. 2020;69(1):147–157.

78. Zheng L, Wang J, Jin X, et al. Erythroblastic island: the niche for erythroid terminal differentiation and beyond. Blood Sci. 2025;7(2):e00228.

79. Yapp C, Nirmal AJ, Zhou F, et al. Highly multiplexed 3D profiling of cell states and immune niches in human tumors. Nat Methods. 2025;22(10):2180–2193.

