## Supplemental Materials for "Niche-level immune evasion in *TP53* mutant AML residual disease revealed by spatial proteomics"

#### Supplemental Methods

##### Patients and samples

From MD Anderson's Institutional Tissue Bank, we obtained archival formalin-fixed, paraffin-embedded (FFPE) bone marrow (BM) biopsy specimens collected at diagnosis and at morphological complete remission (CR; at the end of induction therapy and/or immediately before the next treatment) from the ilia of acute myeloid leukemia (AML) patients diagnosed or treated at MD Anderson between 2016 and 2023. All specimens had been processed according to the standard laboratory protocol in a CLIA-certified clinical pathology laboratory. To enable immunofluorescence-based detection of leukemia cells, *TP53* mutant samples with missense mutations and *TP53* wild-type samples with *NPM1* mutations were selected. We also obtained control normal BM (NBM) specimens collected from patients who were initially suspected of having hematologic disease but for whom pathological assessment revealed no abnormal histological findings, no gene mutations characteristic of hematologic disorders or cytogenetic abnormalities. Patients with monoclonal gammopathy of undetermined significance or lymphoma without BM involvement were not included. AML patients were enrolled in clinical trials as part of MD Anderson's AML "Moonshot" program and Break Through Cancer's "Eradicating MRD in AML" project, and sample collection and analysis were performed under protocols approved by MD Anderson's Institutional Review Board (PA19-0800/LAB02-395). Written informed consent was obtained from all patients in accordance with the Declaration of Helsinki. Each patient's *TP53* mutation status at diagnosis was determined according to previously described criteria.[1]

##### seqIF staining and imaging

Slides for seqIF were prepared in MD Anderson's Research Histology Core Laboratory. FFPE tissue samples were cut into 5-μm sections, which were placed on slides and subjected to preprocessing. The slides were incubated at 60°C overnight followed by deparaffinization and rehydration, consisting of washes in xylene (3×5 minutes), 100% ethanol (2×3 minutes), 70% ethanol (2×3 minutes), 50% ethanol (2×3 minutes), 30% ethanol (2×3 minutes), and phosphate-buffered saline (PBS; 1×2 minutes). The slides were then incubated in 10% hydrogen peroxide with PBS for 30 minutes and subjected to heat-induced antigen retrieval at 107°C for 15 minutes in EZ-AR2 buffer using an EZ-Retriever microwave (both from BioGenex). After cooling to room temperature, the slides were incubated in quenching buffer (Lunaphore Technologies) for 10 minutes and stored in PBS. The slides were then loaded into the COMET system (Lunaphore Technologies) for seqIF staining and processed according to the manufacturer's recommendations. The samples underwent staining cycles with primary and secondary antibodies (listed in **Supplemental Table 2**) and were imaged, and then antibody elution was performed as described previously.[2] The incubation times for the primary antibodies (optimized in each antibody cycle) were set at 4 or 8 minutes, and those for the secondary antibodies were set at 2 minutes. All fluorescence signals were detected in the Cy5 channel, except that of αSMA, which was detected in the TRITC channel. The output image files were multi-layer ome.tiff images comprising DAPI and autofluorescence images obtained before the staining cycles and automatically layered images obtained during each cycle.

**Data analysis**

**Cell segmentation and classification**

Images were loaded into Visiopharm software. First of all, regions of interest (ROIs) were set as a whole bone marrow area. Initial quality filtering comprised the detection of tissue and the removal of artifacts such as tissue folding, out-of-focus areas, misalignment across obtained images, broken tissues, extremely high autofluorescence areas, weak DAPI staining, and antibody aggregates. Areas with commonly positive signals in multiple unrelated layers, such as FOXP3, p53, CD71, CD8, and CD105 signals, were also removed. After artifact removal, ROIs were further trimmed to exclude their edges, as peripheral regions tend to introduce noise into spatial analysis. Nuclei detection was performed with a pretrained deep learning–based nuclei detection application in Visiopharm using DAPI staining images. Nuclear regions less than 6 $\mu m^2$ were removed after nuclei detection. Borders were drawn along the midlines of the edges of two neighboring nuclei or 5 μm from the edges of detected nuclei regions. The nuclei regions were expanded either 3 μm or until they reached the drawn borders and were defined as cell regions. Cell segmentation in the images was manually reviewed and modified if needed. The pixel intensities of each layer of the fluorescence images were subtracted, pixel by pixel, from the corresponding pixel intensities of the matched autofluorescence image captured in the first cycle, to remove autofluorescence from the original image, and the resulting images were filtered using a 3×3-pixel median filter to reduce signal noises. For p53+ cell detection, the mean signal intensity of each nuclear region was compared to a threshold value to determine p53 positivity. The threshold value was selected to minimize p53+ cell detection in *TP53* wild-type ($TP53^{WT}$) BM at CR and in NBM. For cell classification, the pixels within each cell region were ranked by signal intensity, and the mean signal intensity of the pixels within the optimized percentile range was compared to a threshold value optimized for each antibody and sample. For cell membrane and cytoplasmic markers, the range was from the 70th-80th percentile to the 95th percentile, whereas for nuclear markers, the range was from the 50th percentile to the 95th percentile, to minimize misclassification due to overlapping cells. In rare cases in which residual antibodies from previous cycles caused a high signal, the affected pixels were removed from the layer prior to the classification step. In cases in which the deep learning–based cell segmentation with DAPI staining images did not align with the cell membrane staining and caused misclassifications, the cell region boundary was manually modified. For larger cells such as myeloblasts, the cell regions were expanded an additional 2 μm from the drawn boundary before classification. Areas with positive CD36 staining, negative CD45 staining, and at least one nucleus with a rich cytoplasm region were manually segmented and classified as megakaryocytes. NK-cell annotation was manually performed based on CD56 and CD45 positivity. Classified cells were manually reviewed and reclassified if necessary. The markers used for each cell type are listed separately (**Supplemental Table 3, Supplemental Figure 4B**). For each sample, the x- and y-coordinates of each cell's center position, along with the cell's classification and marker intensities, were exported from Visiopharm into a CSV file.

**Structure segmentation and classification**

In an initial step, structures such as MSCs, endothelial cells (ECs), adipocytes, and trabecular bone were segmented and classified using thresholds of corresponding marker expressions and modifications based on morphological features optimized for each sample, pixel by pixel. The markers used for classification are listed separately (**Supplemental Table 3**). Modifications based on morphological features included the removal of miniscule regions and the removal of low-circularity regions for adipocytes. All regions were manually reviewed and refined using the marker combinations used in the initial step. Adipocytes were initially identified as empty areas and subsequently annotated as adipocytes if they exhibited a round morphology or were delineated by surrounding CD36 staining in the absence of MSCs or ECs. In cases with an incomplete adipocyte boundary, the two visible ends of the CD36+ rim were manually connected to approximate the full contour. In cases with discontinuous, fragmented, or shifted CD56 staining of trabecular bone, the putative structural boundaries of the trabecular bone were delineated based on the morphological features in several fluorescence images including autofluorescence images. Arterioles were manually annotated based on morphologically distinct regions exhibiting high CD34 and αSMA expression. Structures immediately outside the ROI were also annotated to ensure accurate assessment of the space between cells and surrounding structures. For each sample, the resulting structure segmentation lines and classifications were exported from Visiopharm into an XML file.

**Protein expression normalization**

Signal intensities were calculated by subtracting the autofluorescence intensity from the raw intensity on a pixel-by-pixel basis and then filtered using a 3×3-pixel median filter to remove signal noises. For each cell region, the mean signal intensity was computed using pixels within the 70th-95th or 80th-95th percentile range for cell membrane and cytoplasmic markers and pixels within the 50th-95th percentile range for nuclear markers. Each samples' mean signal intensities were normalized using Z-score normalization and then used for heatmap construction and visualization.[3] To compare protein expression across samples, we used the UniFORM-based method implemented in Python (3.9.6) to normalize signal intensities.[4] Since our dataset contained many near-zero values and integer-valued measurements for FOXP3 and TIGIT, we employed arcsinh transformation rather than the log normalization with thresholding used in the original UniFORM pipeline. We applied Gaussian smoothing before constructing the FDataGrid object. Furthermore, to preserve the integrity of the entire dataset, we retained the transformed values beyond the grid ranges after applying the shift function, instead of setting them to zero, as in the original method. The distributions of normalized values are shown separately (**supplemental Figure 19**).

**Normalized median shortest distance analysis**

Cell-cell spatial proximity was quantified by calculating the shortest distances between each cell and the nearest cell of a defined cell class or cluster. For each sample, the cells' x- and y-coordinates and class annotations were extracted from the single-cell segmentation dataset. For each cell type of interest, the Euclidean distances from all cells to their nearest neighboring cell of that cell type were computed. In calculating the distance from one cell to the nearest cell the same type, the second-nearest neighbor distance was used to avoid self-matching. Next, distances were aggregated by cell type. For each cell type pair, the median shortest distance was calculated, providing a robust measure of spatial proximity. To enable cross-sample comparison, we normalized these medians by dividing each value by the mean distance across all phenotypes within the same sample, resulting in a normalized median shortest distance (NMSD) matrix. To assess the significance of the observed proximities, we generated a permutation-based null model. For each sample, the cell type labels were randomly permuted 1,000 times while the cell coordinates were preserved. For each permutation, the median shortest distances were recalculated as above. Empirical p-values were computed as the proportion of permuted medians smaller than the observed values. To test the spatial segregation of the clusters, we computed p-values as the proportion of permuted medians greater than the observed values. To combine p-values across samples, we applied the Stouffer method, which yielded consensus significance scores for each phenotype pair. To visualize NMSDs, we extracted the distances from all other cell types to a target cell type. The group-wise means and standard deviations (SDs) of the NMSDs were calculated, and significance values were derived from either permutation tests (for single-group analysis) or Wilcoxon rank-sum tests (for two-group comparisons). Meta p-values were computed using the Stouffer method.

**Spatial neighborhood enrichment analysis**

Spatial neighborhood enrichment (SNE) analysis using cell type classifications was performed as described previously.[5,6] Up to predetermined numbers of the nearest neighbor cells within 30 μm of each cell were collected. Each cell cluster was computed based on the similarities of the neighboring cell types using K-means clustering. The K-centroids representing the neighboring cell type composition of each cluster were normalized by the overall frequency of each cell type across the tissues. $Log_2$ fold-changes were calculated to quantify the enrichment or depletion of each cell type relative to the tissue average. A hypergeometric test was performed to assess whether the number of a specific cell type observed within a given cluster was significantly higher than that expected by chance. For each cell type and cluster pair, the p-value represents the probability of observing at least the actual number of that cell type in the cluster, given its overall abundance in the tissue. The resulting p-values were adjusted using the Benjamini-Hochberg method, and statistical significance was defined as an adjusted p-value <0.05.

**Structure proximity analyses**

Structure proximity analyses for both cell types (SPA-cell) and clusters (SPA-cluster) were performed as described previously[5] with slight modifications optimized for our dataset. Briefly, the shortest distances from each cell grouped by class to the nearest structures were calculated. The spatial distribution of each class was modeled using a Poisson point process model with the intensity function in spatstat (3.4.1) in R (4.5.1):[7]

$$\lambda(u) = \exp(\beta_0 + \beta_1 S(u))$$

where $u$ represents a spatial location in the tissue, $S(u)$ is the distance from each structure, $\beta_0$ represents the baseline cell density near the structure, and $\beta_1$ describes how the density changes with increasing distance from the structure. Then, we ranked the cell classes' $-\beta_1$ values and scaled the ranking number with min-max normalization in each sample. For the significance test, cell classification labels were randomly permutated 100 times. In each permutation, the median shortest distance between each cell class and each structure was recalculated. The p-value was defined as the proportion of permutations in which the median distance was shorter than the actual observed median distance, indicating whether the observed proximity was greater than that expected by chance. To combine the p-values across samples, we calculated meta p-values using the Stouffer method.

**Protein expression and distance analysis**

Euclidean distances between two cells were calculated using the x- and y-coordinates of each cell's center. The spatstat package was used to calculate distances between cells and structures using point pattern objects created from the x- and y-coordinates of the cells' centers and line segment objects derived from the structure segmentation masks in XML files. Cells were grouped according to their proximity to a target cell or structure class, defined by the shortest distances, and the normalized expression of proteins were compared between these proximity-based groups. Significance was determined using the Wilcoxon rank-sum test.

**Visualization of cells and structures**

Cells and structures were visualized using the spatstat package. Original CSV and XML files exported from Visiopharm were converted to ppp objects and psp objects, respectively, and plotted. Background gradation was drawn using the distmap function in spatstat[7].

**Cluster enrichment near individual MSC structures**

To identify the MSC structure closest to each cell, we computed the perpendicular Euclidean distance from every cell to all MSC boundary segments. Connected line-segment objects were then grouped into individual MSC structural units using the igraph package to ensure structural continuity. Each cell was assigned to the MSC component corresponding to its nearest segment. Cells belonging to leukemia cell clusters (p53/p53CD34/p53CD71-1 and -2) and lymphocyte clusters (T-1 and T-2) were extracted for analysis. MSC components with fewer than five cells were excluded, as were cells more than 50 μm from any MSC component. Differences in the numbers of leukemia cells and lymphocytes associated with MSC components were assessed using chi-square tests of independence; p-values were obtained by permutation to account for low expected counts. For MSC components that passed the filtering criteria, each structure was classified as leukemia-enriched (>75% leukemia), lymphocyte-enriched (<25% leukemia), or mixed (25-75% leukemia), and the distribution of these categories for each sample was visualized.

**scRNA-seq data acquisition and cell-cell communication analysis**

After informed consent was obtained, samples were collected according to institutional guidelines under IRB-approved protocols (BTC/PA19-0800/LAB02-395). BM aspirates collected from leukemia patients were subjected to mononuclear cell isolation and scRNA-seq performed using the Chromium single-cell platform (10x Genomics). The Chromium Next GEM Single Cell 3' GEM v3.1 kit was used for scRNA-seq library preparation. Initial data processing (up to the generation of the count matrix of the obtained data) was performed using Cell Ranger software[8] (10x Genomics) according to the manufacturer's recommendations. The details of the initial data processing step were described previously.[9] Briefly, output files from Cell Ranger were subjected to data quality check, batch correction, UMAP dimension reduction, clustering, and cell annotation using Seurat v5[10] and BoneMarrowMap[11] with the standard data processing protocol. Differentially expressed genes were identified using the FindMarkers function with a default 'test.use' argument in Seurat v5. Cell-cell communication analysis was performed using the R package CellChat v2[12] with CellChatDB, which includes the GDF15-CD48 interaction.[13]

**Data and code availability**

This paper does not report original code. The "van Galen 2019" and "Zeng 2025" scRNA-seq datasets reanalyzed in this study are found in GSE116256 and GSE289435, respectively.[11,14] Any additional information required to reanalyze the data reported in this paper is available from the corresponding author upon request.

#### References


1. Arber DA, Orazi A, Hasserjian RP, et al. International Consensus Classification of Myeloid Neoplasms and Acute Leukemias: integrating morphologic, clinical, and genomic data. *Blood*. 2022;140(11):1200-1228.
2. Rivest F, Eroglu D, Pelz B, et al. Fully automated sequential immunofluorescence (seqIF) for hyperplex spatial proteomics. *Sci Rep*. 2023;13(1):16994.
3. Hickey JW, Tan Y, Nolan GP, Goltsev Y. Strategies for Accurate Cell Type Identification in CODEX Multiplexed Imaging Data. *Front Immunol*. 2021;12:727626.
4. Wang K, Ait-Ahmad K, Kupp S, et al. Toward universal immunofluorescence normalization for multiplex tissue imaging with UniFORM. *Cell Rep Methods*. 2025;5(9):101172.
5. Bandyopadhyay S, Duffy MP, Ahn KJ, et al. Mapping the cellular biogeography of human bone marrow niches using single-cell transcriptomics and proteomic imaging. *Cell*. 2024;187(12):3120-3140.e3129.
6. Schürch CM, Bhate SS, Barlow GL, et al. Coordinated Cellular Neighborhoods Orchestrate Antitumoral Immunity at the Colorectal Cancer Invasive Front. *Cell*. 2020;182(5):1341-1359.e1319.
7. Baddeley A, Turner R. spatstat: An R Package for Analyzing Spatial Point Patterns. *Journal of Statistical Software*. 2005;12(6):1 - 42.
8. Zheng GX, Terry JM, Belgrader P, et al. Massively parallel digital transcriptional profiling of single cells. *Nat Commun*. 2017;8:14049.
9. Li L, Muftuoglu M, Ayoub E, et al. Somatic TP53 Mutations Drive T and NK Cell Dysfunction in AML and Can be Rescued by Reactivating Wild Type p53. *medRxiv*. 2025:2025.2004.2011.25325281.
10. Hao Y, Stuart T, Kowalski MH, et al. Dictionary learning for integrative, multimodal and scalable single-cell analysis. *Nat Biotechnol*. 2024;42(2):293-304.
11. Zeng AGX, Iacobucci I, Shah S, et al. Single-cell Transcriptional Atlas of Human Hematopoiesis Reveals Genetic and Hierarchy-Based Determinants of Aberrant AML Differentiation. *Blood Cancer Discov*. 2025;6(4):307-324.
12. Jin S, Plikus MV, Nie Q. CellChat for systematic analysis of cell-cell communication from single-cell transcriptomics. *Nat Protoc*. 2025;20(1):180-219.
13. Wang Z, He L, Li W, et al. GDF15 induces immunosuppression via CD48 on regulatory T cells in hepatocellular carcinoma. *J Immunother Cancer*. 2021;9(9):e002787.
14. van Galen P, Hovestadt V, Wadsworth Ii MH, et al. Single-Cell RNA-Seq Reveals AML Hierarchies Relevant to Disease Progression and Immunity. *Cell*. 2019;176(6):1265-1281 e1224.

**Supplemental Table 1**
**Patient characteristics**

| Patient ID | Age | Sex | Blast at diagnosis (%) | FC-MRD (%) | Molecular MRD (%) | Gene mutations | Karyotype | Treatment | TP53 status |
|---|---|---|---|---|---|---|---|---|---|
| 1 | 81 | F | 24% | 0.01% | Yes | TP53 | complex | Aza, Ven, Magrolimab | Multi-hit |
| 2 | 67 | M | 22% | 0.60% | Yes | TP53 | complex | Cladribine, Ven, AraC | Multi-hit |
| 3 | 56 | F | 32% | 0% | Yes | TP53, DNMT3A, IKZF1 | complex | Aza, Ven, Magrolimab | Single-hit |
| 5 | 58 | M | 32% | 0% | Yes | TP53 | complex | Aza, Ven, Magrolimab | Multi-hit |
| 6 | 60 | F | 40% | 3.70% | Yes | TP53 | complex | Cladribine, AraC, IDR | Single-hit |
| 8 | 62 | M | 20% | 0% | Yes | TP53 | complex | DAC, Ven, AraC | Multi-hit |
| 12 | 80 | M | 85% | 0% | No | NPM1, FLT3-ITD, IDH2, SRSF2 | 46,XY | ASTX727, Ven, Gilteritinib | |
| 13 | 72 | F | 95% | 0% | No | NPM1, TET2, SRSF2 | 46,XX | Aza, ven, anti-DR5 | |
| 14 | 62 | F | 87% | 0% | No | NPM1, FLT3-ITD, DNMT3A | 46,XX, del(13)(q12q14) | ASTX727, Ven, Gilteritinib | |
| 15 | 47 | F | 70% | 0% | No | NPM1 | 46,XX | Cladribine, AraC, Ven | |
| 16 | 66 | M | 83% | neg | No | NPM1, IDH1 | 46,XY | aza, ven, ivosidenib -> SCT | |
| 18 | 64 | M | 23% | 1.40% | Yes | TP53, NRAS, EZH2 | complex | Aza, Ven, Magrolimab | Multi-hit |
| 20 | 65 | F | 48% | 3.50% | Yes | TP53, TET2, CALR | complex | DAC, Ruxolitinib | Multi-hit |
| 21 | 66 | F | 35% | 0.20% | No* | TP53, RUNX1 | complex with +8,-21,del3p | Vosaroxin, DAC | no record |
| 24 | 52 | M | 70% | 0.50% | No | NPM1, FLT3-ITD, DNMT3A | 46,XY | CLIA, Ven, midostaurin | |
| 25 | 75 | M | 24% | 0.40% | Yes | TP53, U2AF1 | complex | Aza, Ven, Magrolimab | Multi-hit |
| 26 | 77 | F | 59% | 1.23% | Yes | TP53, NRAS | +8, complex | anti-CD47, Aza | Multi-hit |
| 27 | 47 | F | 60% | 0% | No | NPM1, KRAS, DNMT3A | 46,XX | CLIA, Ven | |
| 29 | 58 | M | 55% | 1.35% | No* | TP53 | complex | Idarubicin, AraC -> Decitabine | no record |

*PCR-based

**Supplemental Table 2**

**Antibodies**

| Name | Source | Clone | Cat# | Dilutions |
|---|---|---|---|---|
| Anti-CD105 antibody [EPR10145-12] | Abcam | EPR10145-12 | ab169545 | 1:50 |
| p53 (DO7) Mouse Monoclonal Antibody | Cell Marque | DO7 | 453M | 1:100 |
| NPM1 (mutant) Polyclonal Antibody | Invitrogen | Polyclonal | PAI1-46356 | 1:200 |
| Anti-CD90 / Thy1 antibody [EPR3133] | Abcam | EPR3133 | ab133350 | 1:50 |
| Rabbit anti-FOXP3 Recombinant Monoclonal Antibody [BLR034F] | Bethyl | BLR034F | A700-034 | 1:100 |
| Mouse anti-CD20 Monoclonal Antibody Purified [L26] | Bethyl | L26 | A500-017ACF | 1:250 |
| Rabbit anti-CD34 Recombinant Monoclonal Antibody [BLR197J] | Bethyl | BLR197J | A700-197CF | 1:1000 |
| Rabbit anti-CD3E Recombinant Monoclonal Antibody [BL-298-5D12] | Bethyl | BL-298-5D12 | A700-016CF | 1:500 |
| Rabbit anti-CD4 Recombinant Monoclonal Antibody [BL-155-1C11] | Bethyl | BL-155-1C11 | A700-015CF | 1:200 |
| Purified anti-human CD8a Antibody | Biolegend | C8/144B | 372902 | 1:100 |
| Rabbit anti-CD14 Recombinant Monoclonal Antibody [BLR187J] | Bethyl | BLR187J | A700-187CF | 1:500 |
| Rabbit anti-CD16 Recombinant Monoclonal Antibody [BLR163J] | Bethyl | BLR163J | A700-163CF | 1:1200 |
| Rabbit anti-CD33 Recombinant Monoclonal Antibody [BLR061G] | Bethyl | BLR061G | A700-061CF | 1:100 |
| α-Smooth Muscle Actin (1A4) Mouse mAb (BSA and Azide Free) | Cell Signaling Technology | 1A4 | 69319 | 1:3000 |
| Anti-TIGIT antibody [BLR047F] - BSA free | Abcam | BLR047F | ab243903-1001 | 1:200 |
| Rabbit anti-NCAM-1/CD56 Recombinant Monoclonal Antibody [BLR152J] | Bethyl | BLR152J | A700-152CF | 1:1250 |
| CD71 (MRQ-48) Mouse Monoclonal Antibody | Cell Marque | MRQ-48 | 171M-9 | 1:100 |
| Rabbit anti-CD36 Recombinant Monoclonal Antibody [BLR238K] | Bethyl | BLR238K | A700-238CF | 1:400 |
| Anti-TGF beta 1 antibody [EPR21143] | Abcam | EPR21143 | EPR21143 | 1:50 |
| Phospho-SMAD2 (Ser465/467) (138D4) Rabbit mAb | Cell Signaling Technology | 138D4 | 138D4 | 1:50 |
| Phospho-SMAD3 (Ser423/425) (C25A9) Rabbit mAb | Cell Signaling Technology | C25A9 | C25A9 | 1:50 |
| F(ab')2-Goat anti-Rabbit IgG (H+L) Cross-Adsorbed Secondary Antibody, Alexa Fluor™ 647 | Invitrogen | Polyclonal | A-21246 | |
| F(ab')2-Goat anti-Mouse IgG (H+L) Cross-Adsorbed Secondary Antibody, Alexa Fluor™ 555 | Invitrogen | Polyclonal | A-21425 | |
| F(ab')2-Goat anti-Mouse IgG (H+L) Cross-Adsorbed Secondary Antibody, Alexa Fluor™ Plus 647 | Invitrogen | Polyclonal | A48289 | |

**Supplemental Table 3**

**Markers used for cell and structure classification**

| Class | Marker |
|---|---|
| T-cell | CD3 |
| CD4 T-cell | CD3 CD4 |
| CD8 T-cell | CD3 CD8 |
| Treg | CD3 FOXP3 |
| Monocyte | CD14 |
| Mature Myeloid | CD16 |
| CD34 HSPC/Blast | CD34 CD105(-) |
| CD33 HSPC/Blast | CD33 |
| Erythroblast | CD71 |
| NK-cell | CD56 CD45 |
| NK/T-cell | CD56 CD3 |
| B-cell | CD20 |
| Megakaryocyte | CD36 CD71(-) CD45(-) |
| p53pos Blast | p53 |
| p53pos CD34 Blast | p53 CD34 |
| p53pos CD71 Blast | p53 CD71 |
| p53pos CD14 Blast | p53 CD14 |
| NPM1c Blast | NPM1c |
| Mesenchymal Stromal Cell | CD105 CD71(-) |
| Endothelial Cell | CD105 CD34 CD71(-) |
| Adipocyte | CD36 TRITC(-) |
| Trabecular Bone | CD56 |
| Arteriole | CD34 $\alpha$SMA |

**Supplemental Table 4**

**False positive rate of p53+ cell detection**

| Pt | group | p53+ | total | positive |
|---|---|---|---|---|
| Pt13 | TP53wt AML CR | 1 | 8811 | 0.0113% |
| Pt15 | TP53wt AML CR | 0 | 20865 | 0.0000% |
| Pt17 | TP53wt AML CR | 2 | 26825 | 0.0075% |
| HD1 | Normal | 0 | 7396 | 0.0000% |
| HD2 | Normal | 0 | 12690 | 0.0000% |
| HD3 | Normal | 0 | 19274 | 0.0000% |

**Supplemental Table 5**

**Clinical Smear Result**

| **Classification** | **Normal range** | **NBM1** | **NBM2** | **NBM3** |
|---|---|---|---|---|
| Blast | 0-5 | 1 | 1 | 2 |
| Progranulocyte | 2-8 | 0 | 1 | 1 |
| Myelocyte | 5-20 | 10 | 7 | 9 |
| Metamyelocyte | 13-32 | 12 | 10 | 8 |
| Granulocyte | 7-30 | 32 | 30 | 35 |
| Eosinophil | 0-4 | 3 | 3 | 2 |
| Lymphocyte | 3-17 | 9 | 12 | 15 |
| Plasma cell | 0-2 | 0 | 1 | 2 |
| Monocyte | 0-5 | 2 | 3 | 5 |
| Reticulum Cell | 0-2 | 0 | 0 | 0 |
| Pronormoblast | 1-8 | 1 | 0 | 1 |
| Normoblast (Erythroblast) | 7-32 | 30 | 32 | 20 |
| M:E Ratio | 3-4 | 1.8 | 1.6 | 2.6 |

**Supplemental Table 6**

**GDF15-CD48 communication probability (predicted_CellType_Broad)**

| Cell Type | B | CD4 Memory T | CD8 Memory T | cDC | Cycling Progenitor | Early Erythroid | Early GMP | Early Lymphoid | EoBasoMast Precursor | HSC MPP | Late Erythroid | Late GMP | LMPP | Megakaryocyte Precursor | MEP | Monocyte | Naive T | NK | pDC | Plasma Cell | Pre-B | Pro-B | Pro-Monocyte | Stromal |
|---|---|---|---|---|---|---|---|---|---|---|---|---|---|---|---|---|---|---|---|---|---|---|---|---|
| B | 0 | 0 | 0 | 0 | 0 | 0 | 0 | 0 | 0 | 0 | 0 | 0 | 0 | 0 | 0 | 0 | 0 | 0 | 0 | 0 | 0 | 0 | 0 | 0 |
| CD4 Memory T | 0 | 0 | 0 | 0 | 0 | 0 | 0 | 0 | 0 | 0 | 0 | 0 | 0 | 0 | 0 | 0 | 0 | 0 | 0 | 0 | 0 | 0 | 0 | 0 |
| CD8 Memory T | 0 | 0 | 0 | 0 | 0 | 0 | 0 | 0 | 0 | 0 | 0 | 0 | 0 | 0 | 0 | 0 | 0 | 0 | 0 | 0 | 0 | 0 | 0 | 0 |
| cDC | 0 | 0 | 0 | 0 | 0 | 0 | 0 | 0 | 0 | 0 | 0 | 0 | 0 | 0 | 0 | 0 | 0 | 0 | 0 | 0 | 0 | 0 | 0 | 0 |
| Cycling Progenitor | 0 | 0 | 0 | 0 | 0 | 0 | 0 | 0 | 0 | 0 | 0 | 0 | 0 | 0 | 0 | 0 | 0 | 0 | 0 | 0 | 0 | 0 | 0 | 0 |
| Early Erythroid | 0 | 0 | 0 | 0 | 0 | 0 | 0 | 0 | 0 | 0 | 0 | 0 | 0 | 0 | 0 | 0 | 0 | 0 | 0 | 0 | 0 | 0 | 0 | 0 |
| Early GMP | 0 | 0 | 0 | 0 | 0 | 0 | 0 | 0 | 0 | 0 | 0 | 0 | 0 | 0 | 0 | 0 | 0 | 0 | 0 | 0 | 0 | 0 | 0 | 0 |
| Early Lymphoid | 0 | 0 | 0 | 0 | 0 | 0 | 0 | 0 | 0 | 0 | 0 | 0 | 0 | 0 | 0 | 0 | 0 | 0 | 0 | 0 | 0 | 0 | 0 | 0 |
| EoBasoMast Precursor | 0 | 0 | 0 | 0 | 0 | 0 | 0 | 0 | 0 | 0 | 0 | 0 | 0 | 0 | 0 | 0 | 0 | 0 | 0 | 0 | 0 | 0 | 0 | 0 |
| HSC MPP | 0 | 0 | 0 | 0 | 0 | 0 | 0 | 0 | 0 | 0 | 0 | 0 | 0 | 0 | 0 | 0 | 0 | 0 | 0 | 0 | 0 | 0 | 0 | 0 |
| Late Erythroid | 0.00827662 | 0.009552247 | 0.009437167 | 0.007653455 | 0 | 0 | 0 | 0 | 0 | 0 | 0 | 0 | 0 | 0 | 0 | 0.00812563 | 0.00946719 | 0.00862421 | 0.001144052 | 0.001893613 | 0.001437861 | 0.000445477 | 0.001880545 | 0 |
| Late GMP | 0 | 0 | 0 | 0 | 0 | 0 | 0 | 0 | 0 | 0 | 0 | 0 | 0 | 0 | 0 | 0 | 0 | 0 | 0 | 0 | 0 | 0 | 0 | 0 |
| LMPP | 0 | 0 | 0 | 0 | 0 | 0 | 0 | 0 | 0 | 0 | 0 | 0 | 0 | 0 | 0 | 0 | 0 | 0 | 0 | 0 | 0 | 0 | 0 | 0 |
| Megakaryocyte Precursor | 0 | 0 | 0 | 0 | 0 | 0 | 0 | 0 | 0 | 0 | 0 | 0 | 0 | 0 | 0 | 0 | 0 | 0 | 0 | 0 | 0 | 0 | 0 | 0 |
| MEP | 0 | 0 | 0 | 0 | 0 | 0 | 0 | 0 | 0 | 0 | 0 | 0 | 0 | 0 | 0 | 0 | 0 | 0 | 0 | 0 | 0 | 0 | 0 | 0 |
| Monocyte | 0 | 0 | 0 | 0 | 0 | 0 | 0 | 0 | 0 | 0 | 0 | 0 | 0 | 0 | 0 | 0 | 0 | 0 | 0 | 0 | 0 | 0 | 0 | 0 |
| Naive T | 0 | 0 | 0 | 0 | 0 | 0 | 0 | 0 | 0 | 0 | 0 | 0 | 0 | 0 | 0 | 0 | 0 | 0 | 0 | 0 | 0 | 0 | 0 | 0 |
| NK | 0 | 0 | 0 | 0 | 0 | 0 | 0 | 0 | 0 | 0 | 0 | 0 | 0 | 0 | 0 | 0 | 0 | 0 | 0 | 0 | 0 | 0 | 0 | 0 |
| pDC | 0 | 0 | 0 | 0 | 0 | 0 | 0 | 0 | 0 | 0 | 0 | 0 | 0 | 0 | 0 | 0 | 0 | 0 | 0 | 0 | 0 | 0 | 0 | 0 |
| Plasma Cell | 0 | 0 | 0 | 0 | 0 | 0 | 0 | 0 | 0 | 0 | 0 | 0 | 0 | 0 | 0 | 0 | 0 | 0 | 0 | 0 | 0 | 0 | 0 | 0 |
| Pre-B | 0 | 0 | 0 | 0 | 0 | 0 | 0 | 0 | 0 | 0 | 0 | 0 | 0 | 0 | 0 | 0 | 0 | 0 | 0 | 0 | 0 | 0 | 0 | 0 |
| Pro-B | 0 | 0 | 0 | 0 | 0 | 0 | 0 | 0 | 0 | 0 | 0 | 0 | 0 | 0 | 0 | 0 | 0 | 0 | 0 | 0 | 0 | 0 | 0 | 0 |
| Pro-Monocyte | 0 | 0 | 0 | 0 | 0 | 0 | 0 | 0 | 0 | 0 | 0 | 0 | 0 | 0 | 0 | 0 | 0 | 0 | 0 | 0 | 0 | 0 | 0 | 0 |
| Stromal | 0 | 0 | 0 | 0 | 0 | 0 | 0 | 0 | 0 | 0 | 0 | 0 | 0 | 0 | 0 | 0 | 0 | 0 | 0 | 0 | 0 | 0 | 0 | 0 |

### Supplemental Figure 1

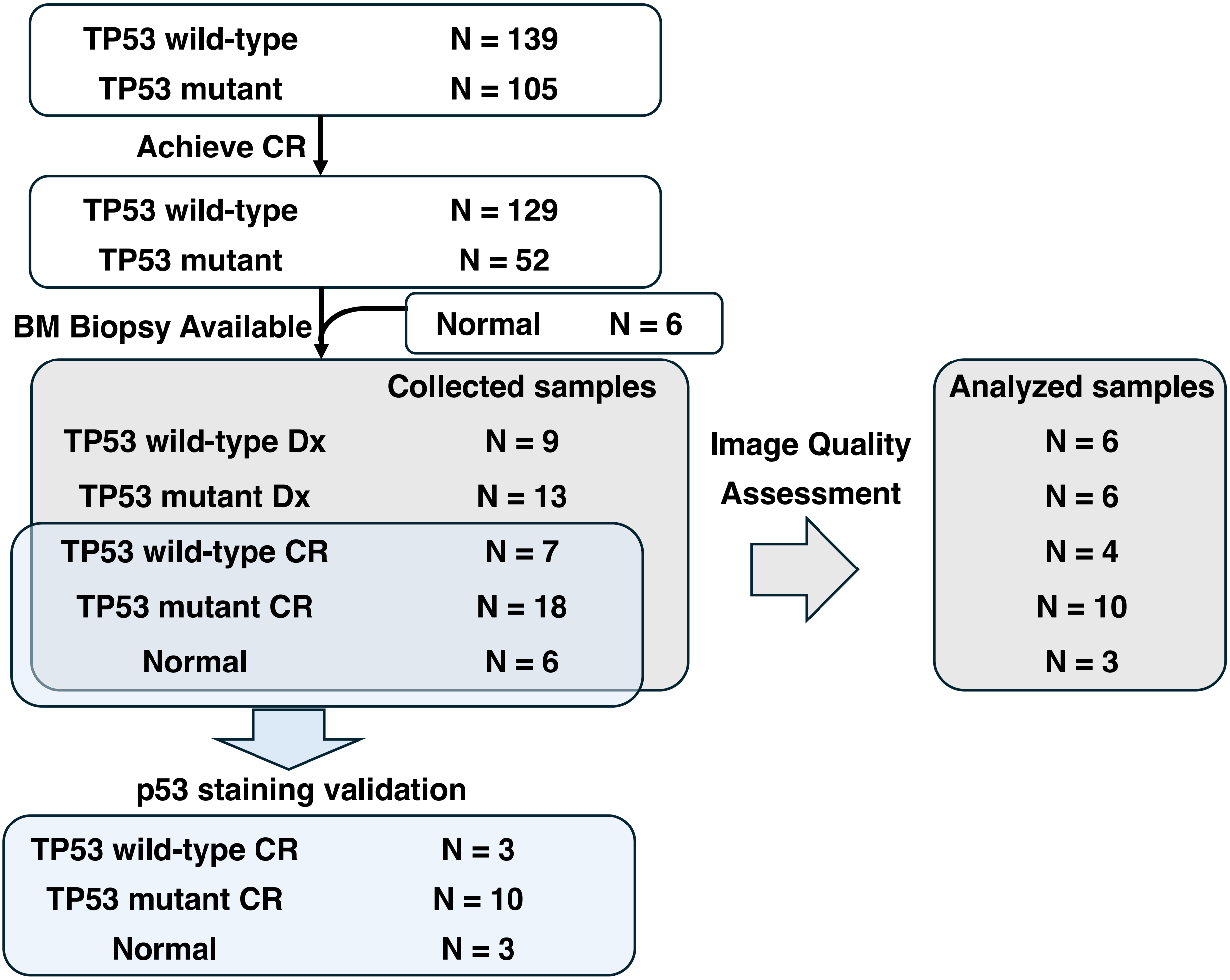

### Supplemental Figure 2

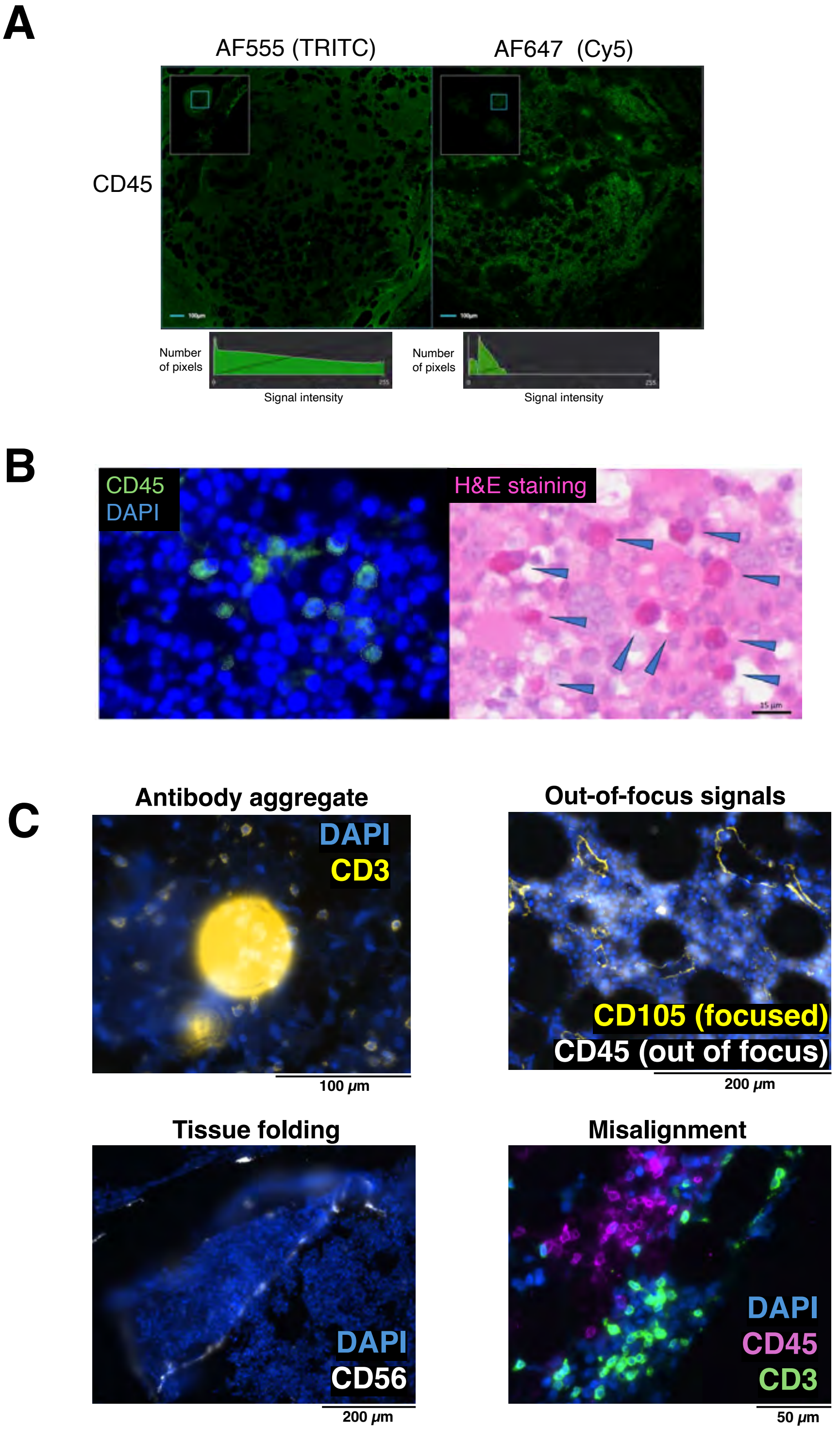

### Supplemental Figure 3

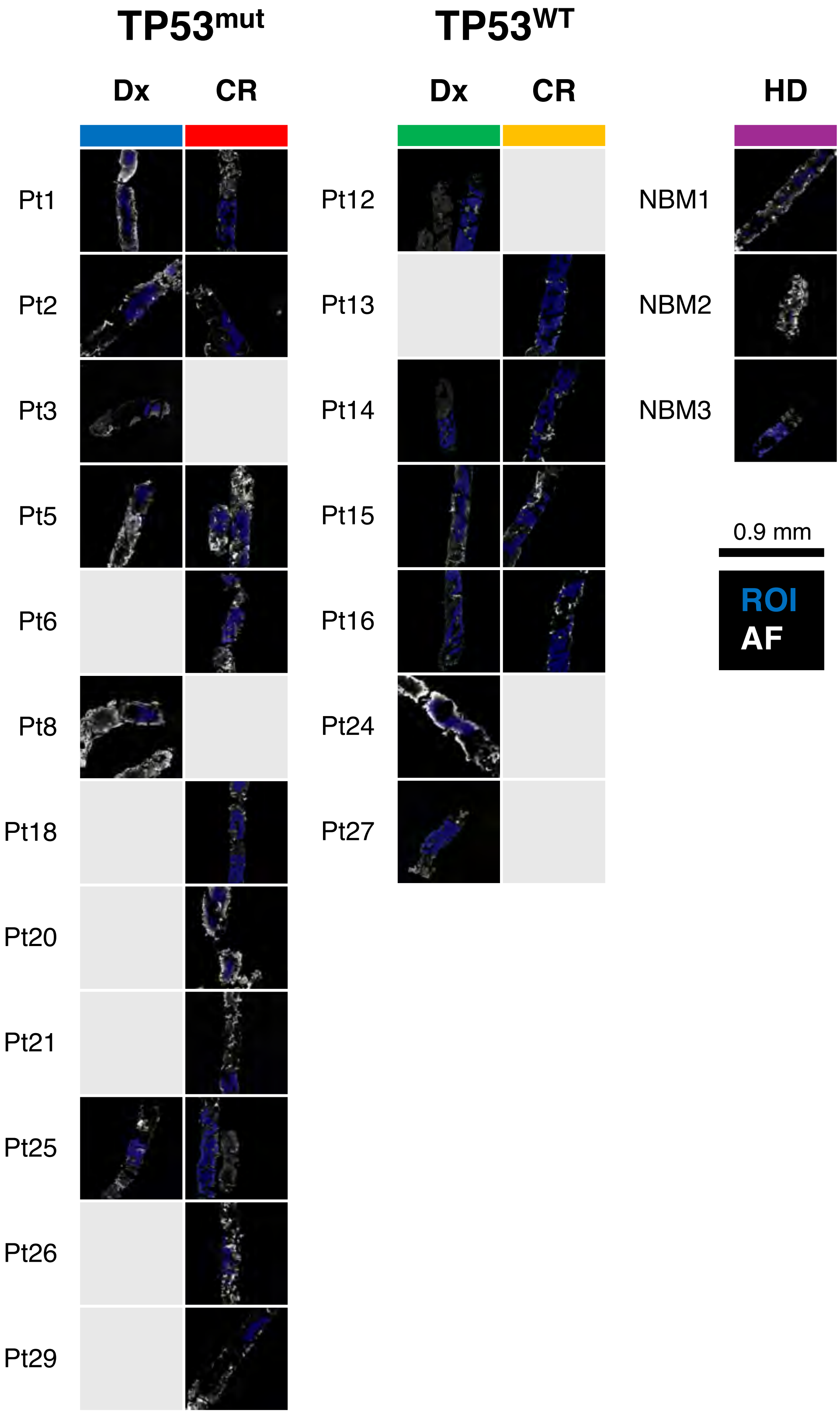

### Supplemental Figure 4

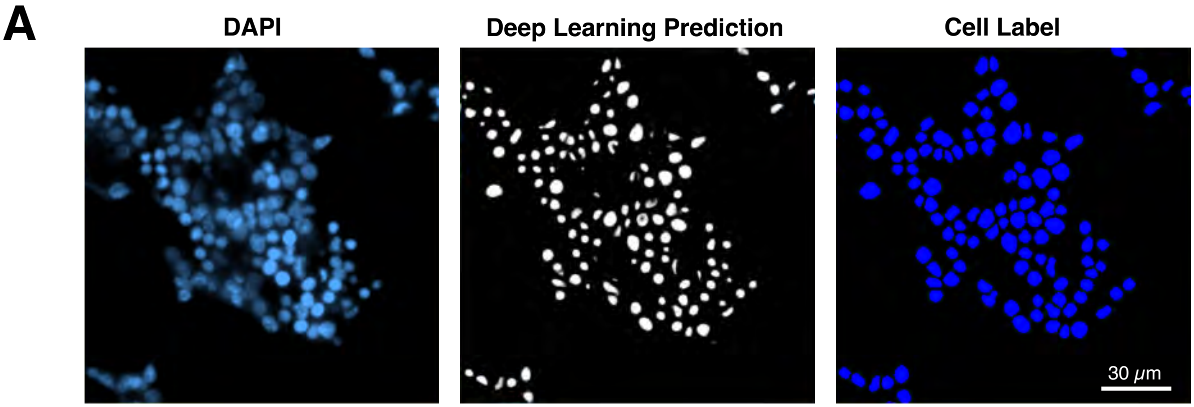


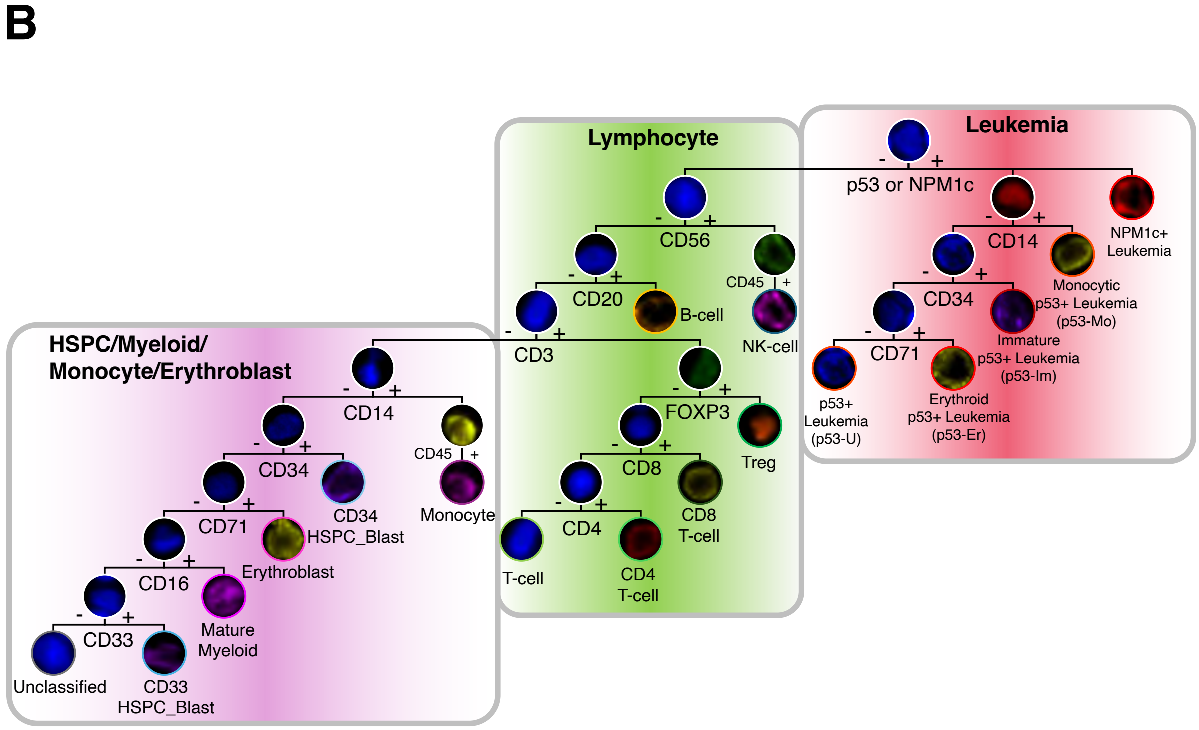

### Supplemental Figure 5

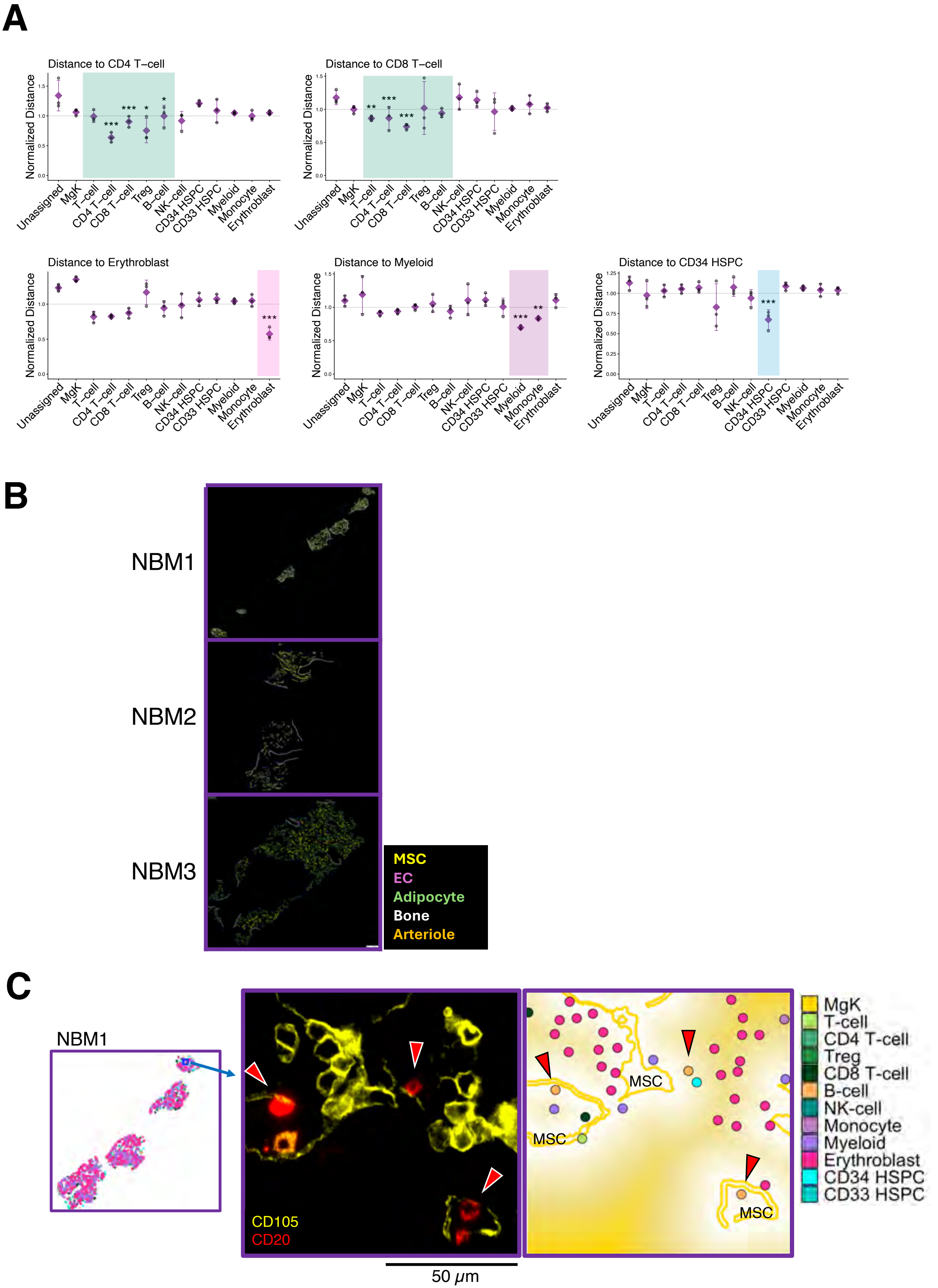

### Supplemental Figure 6

## A

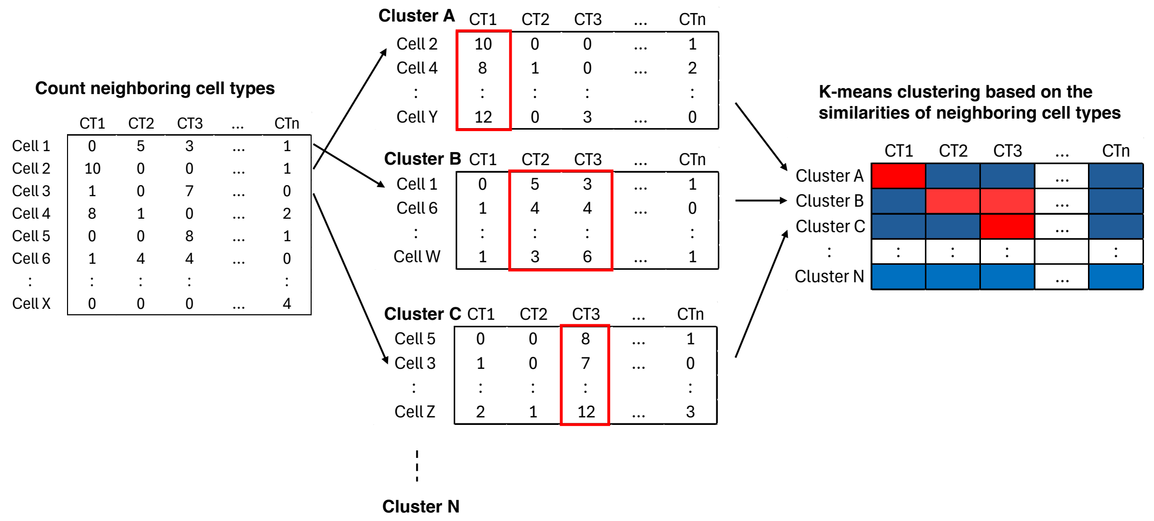


## B

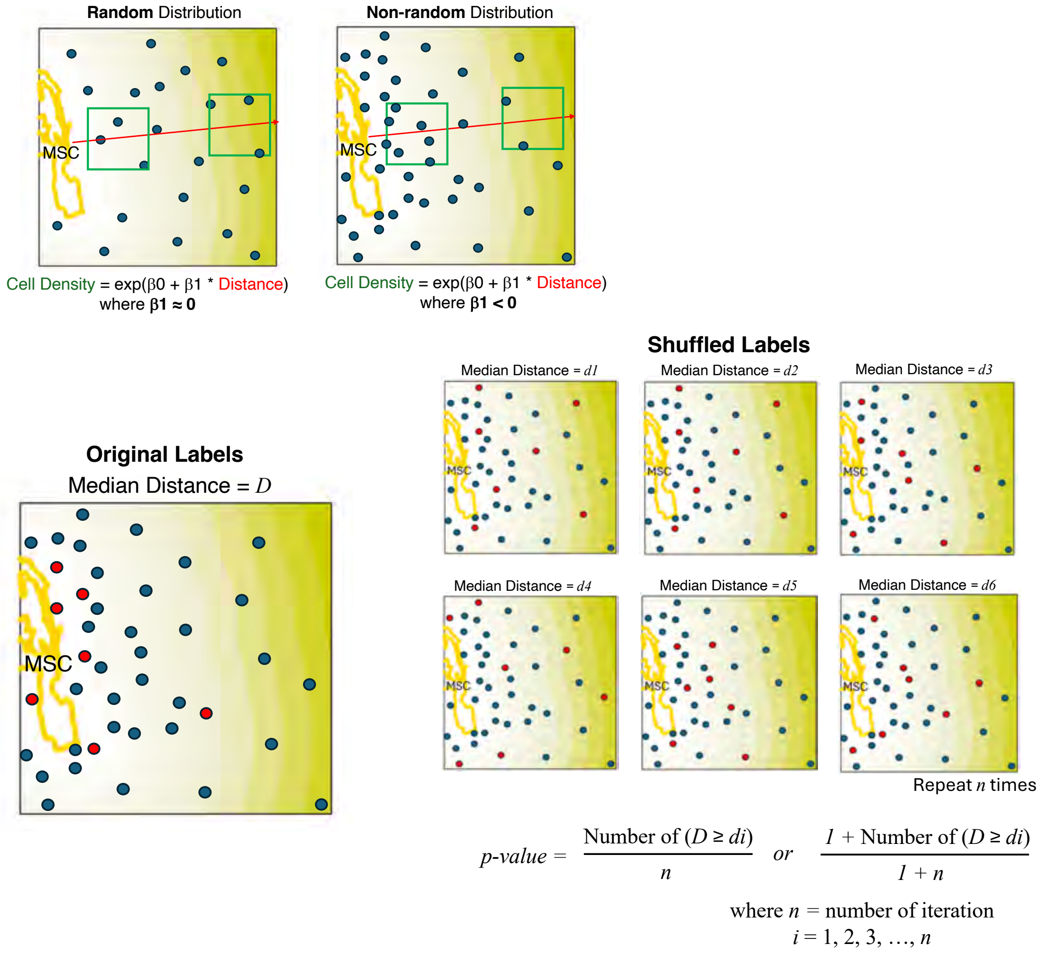

### Supplemental Figure 7

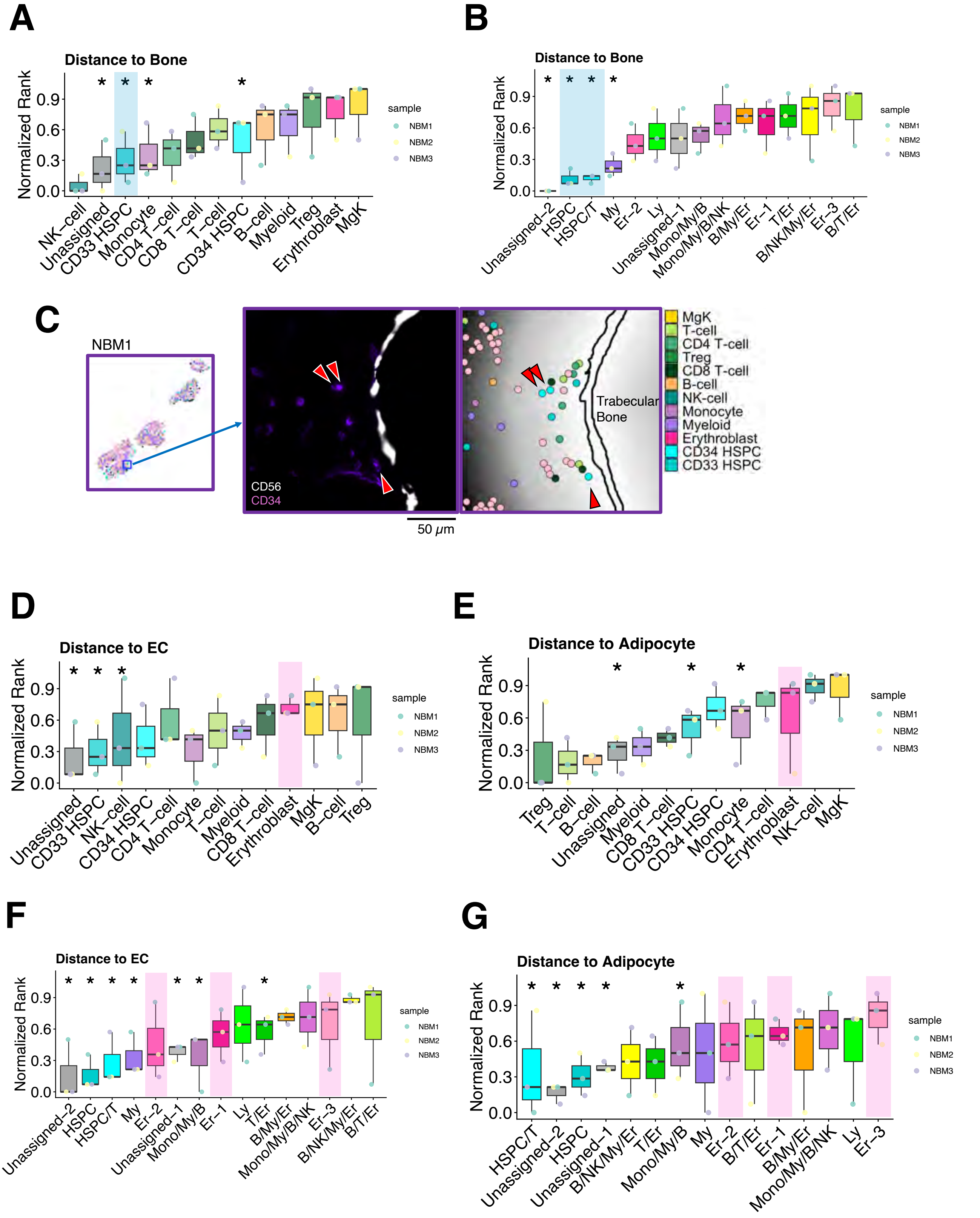

### Supplemental Figure 8

A

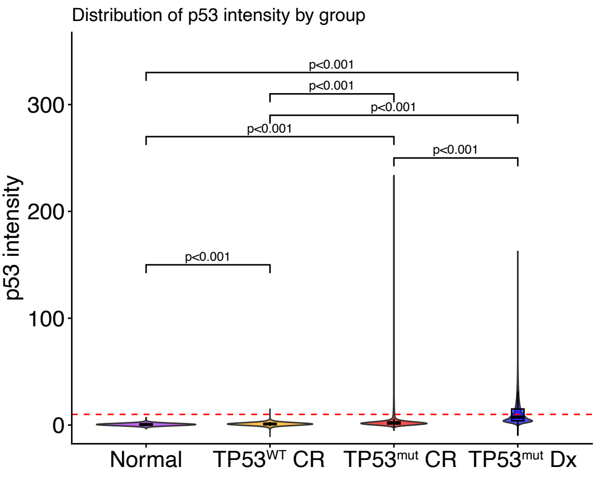


B

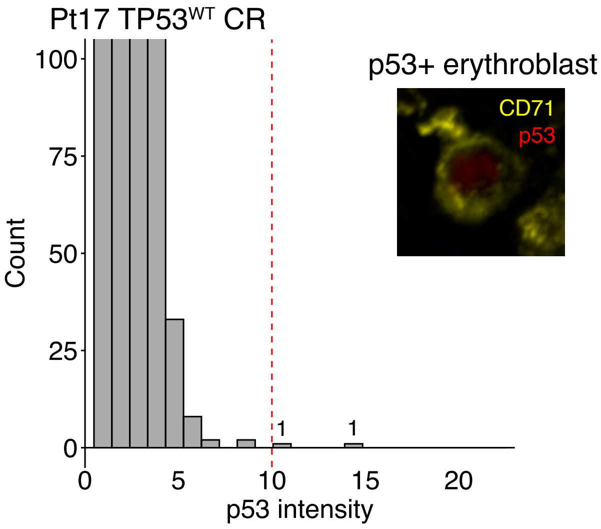


C

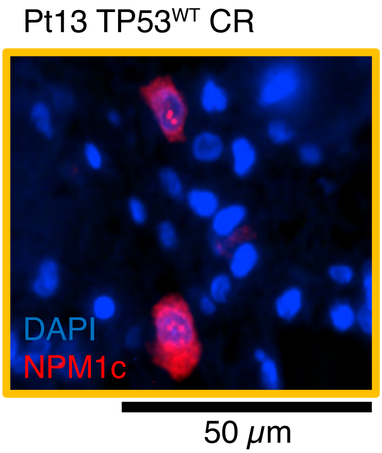

### Supplemental Figure 9

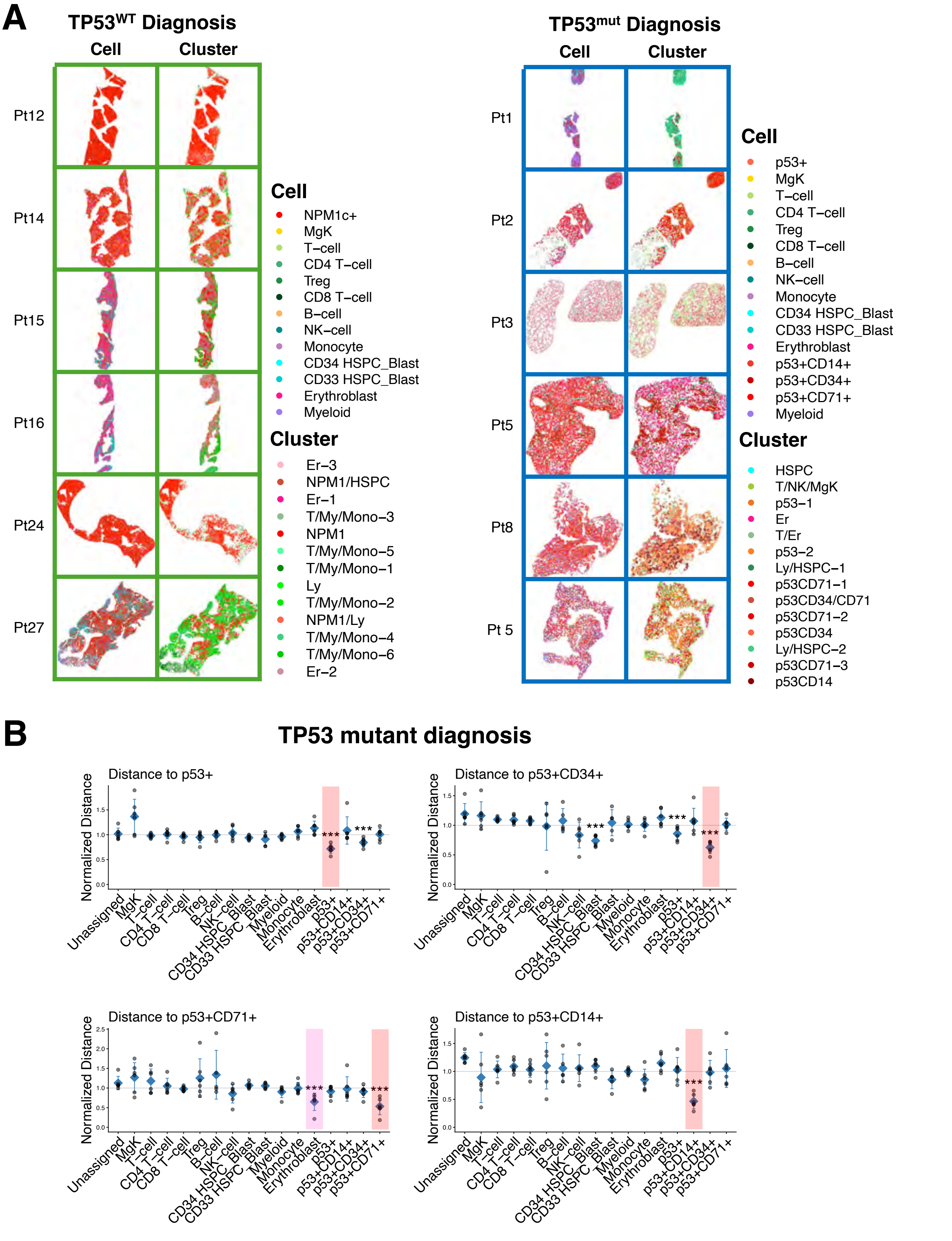

### Supplemental Figure 10

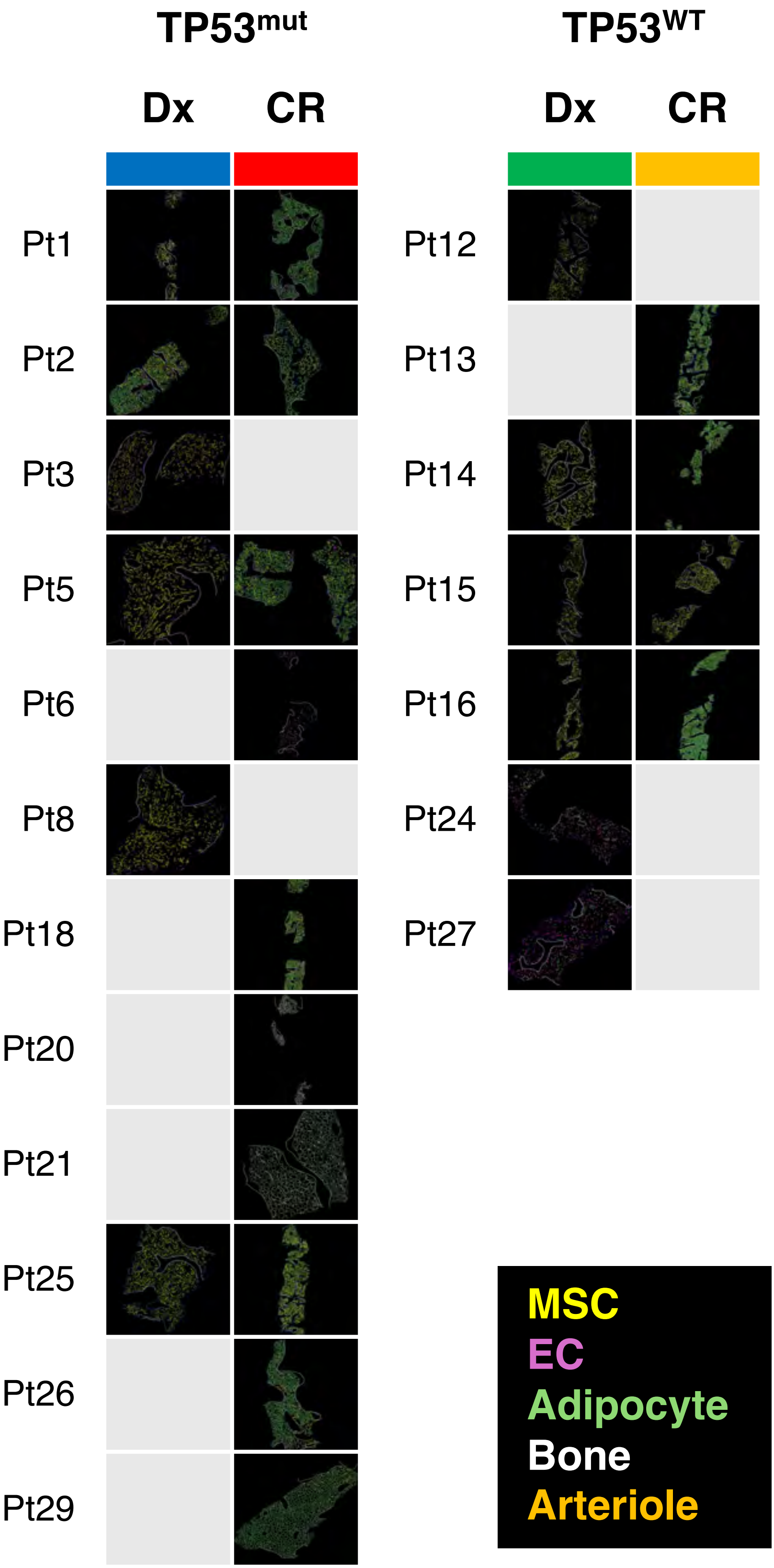

### Supplemental Figure 11

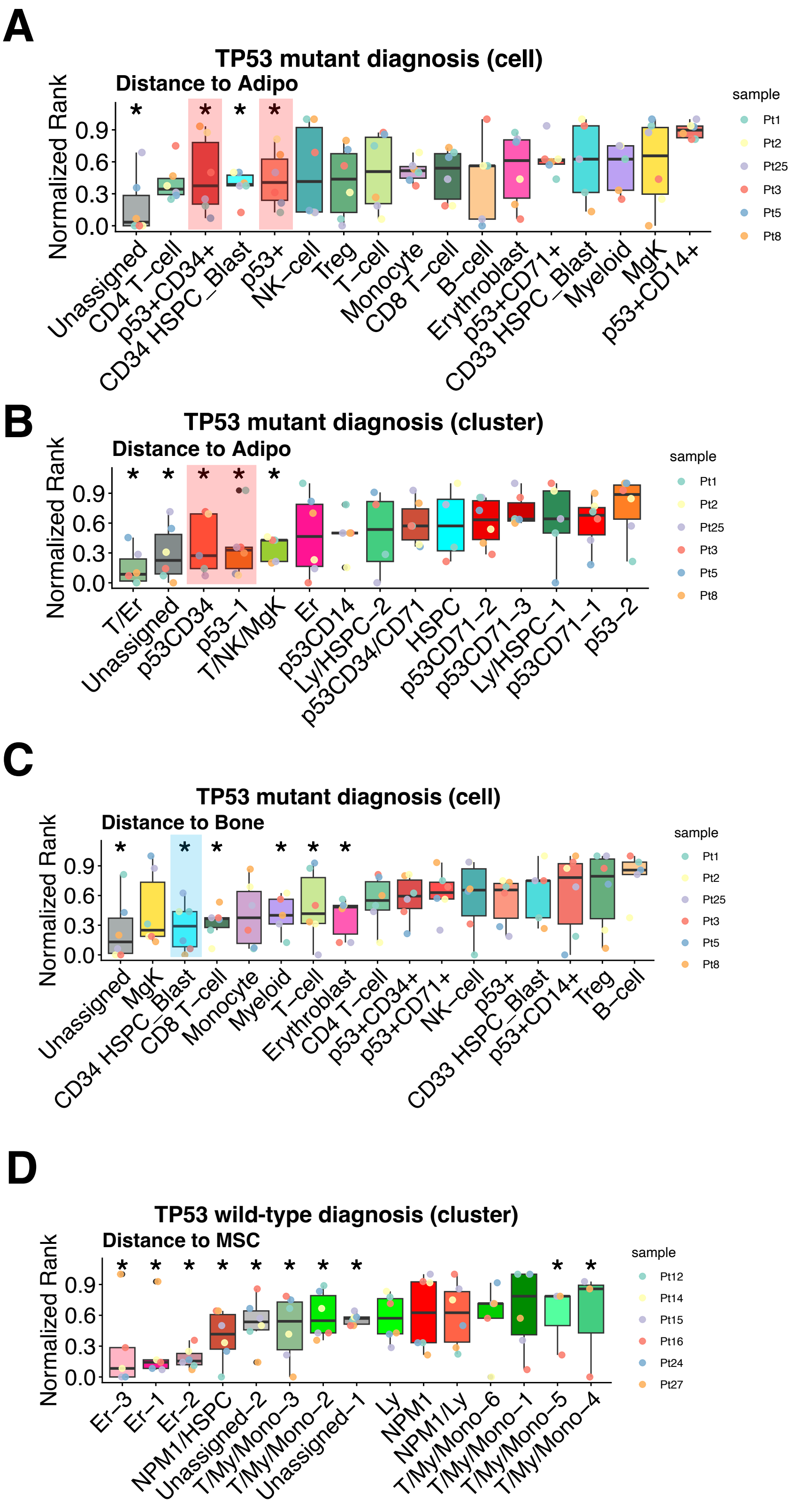

### Supplemental Figure 12

## A

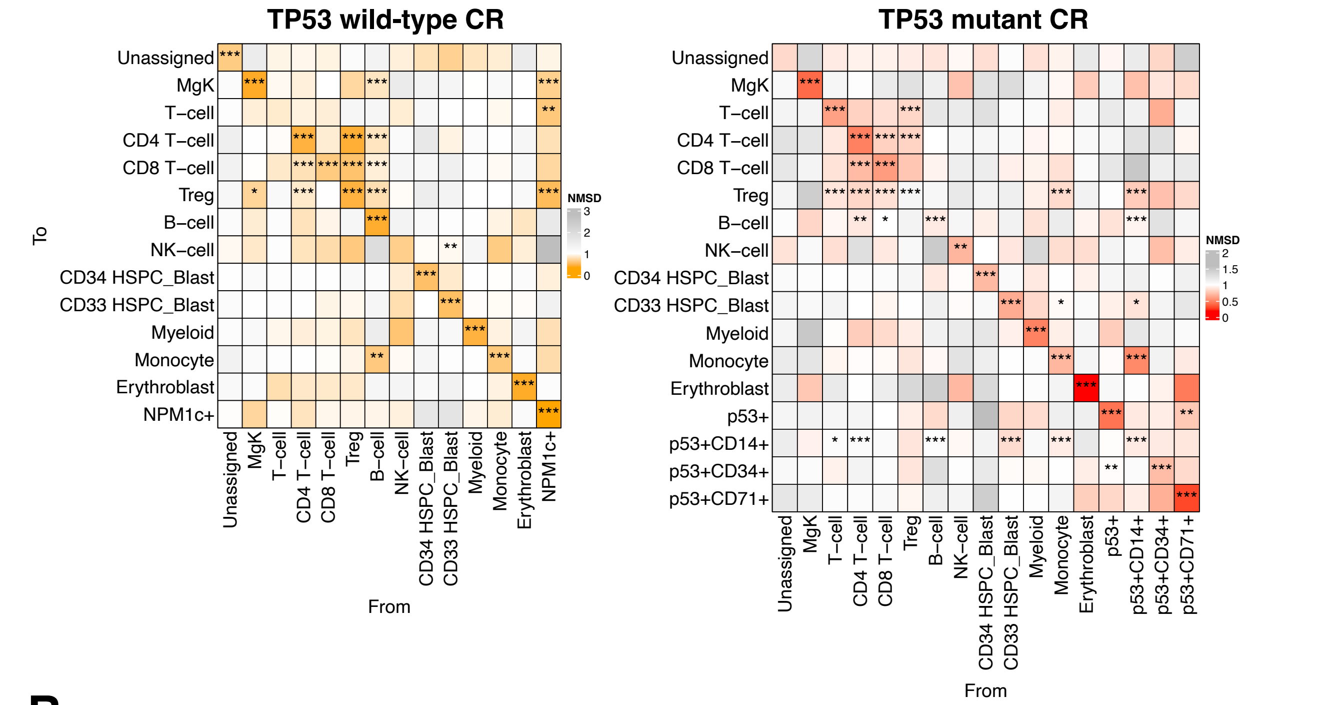


## B

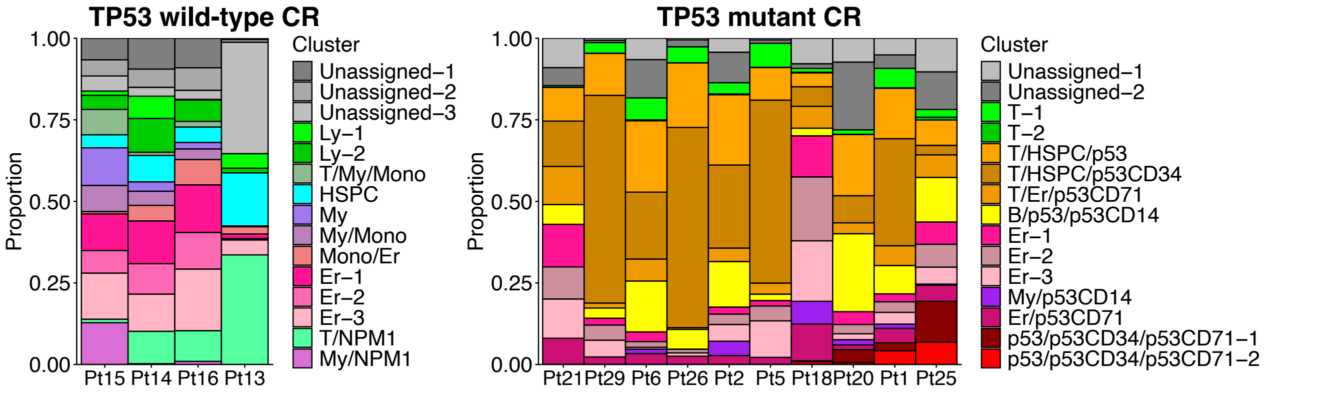

### Supplemental Figure 13

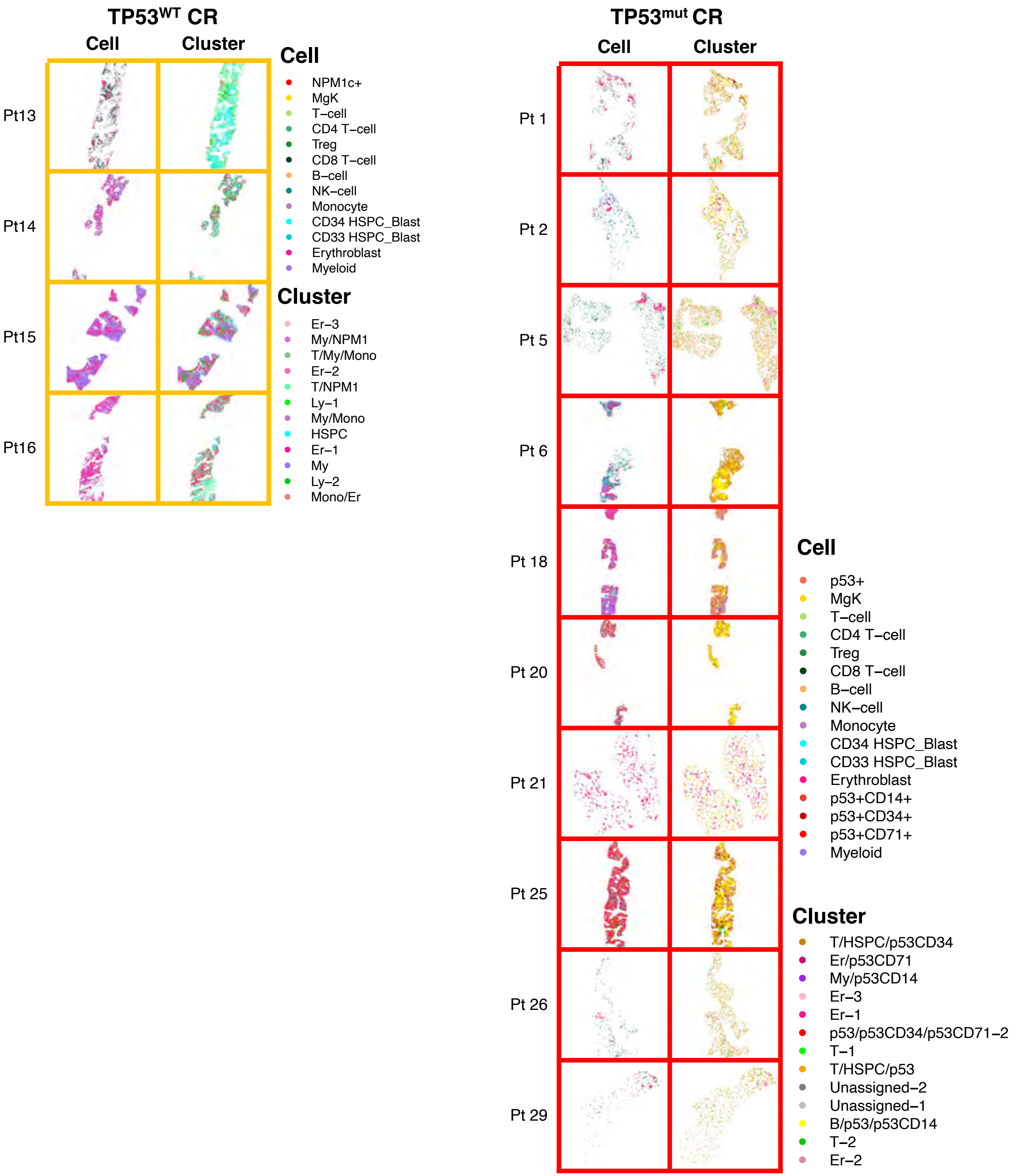

### Supplemental Figure 14

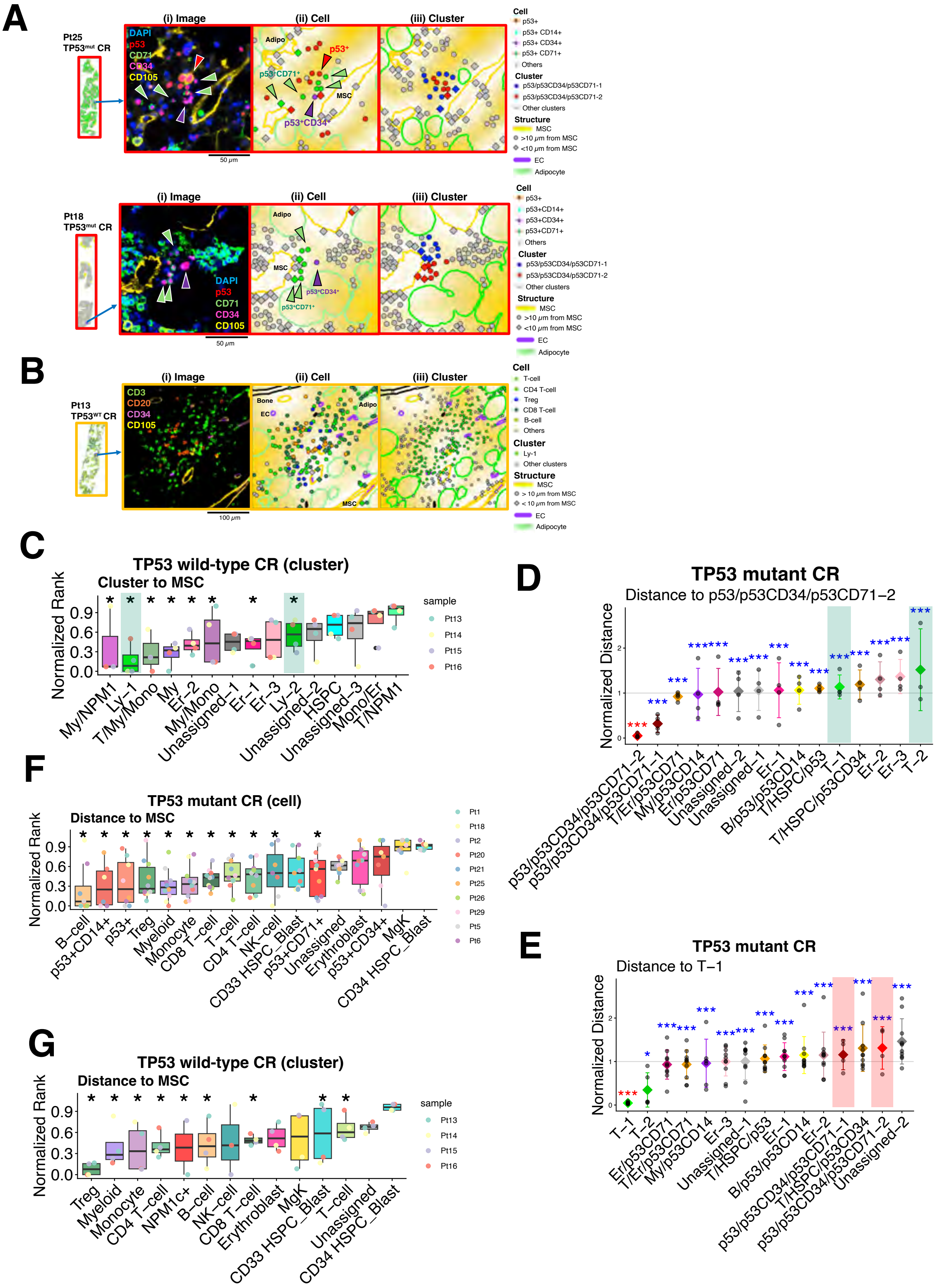

### Supplemental Figure 15

**A**

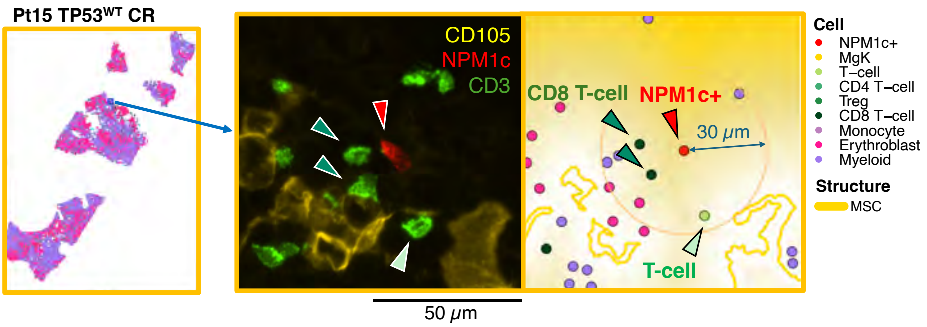


**B**

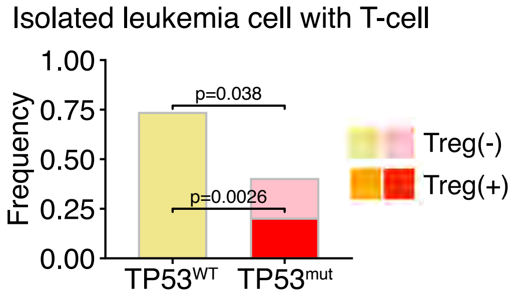


**C**

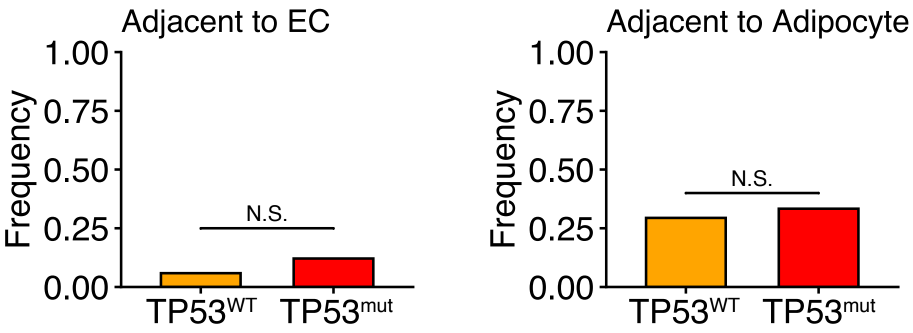

#### Supplemental Figure 16

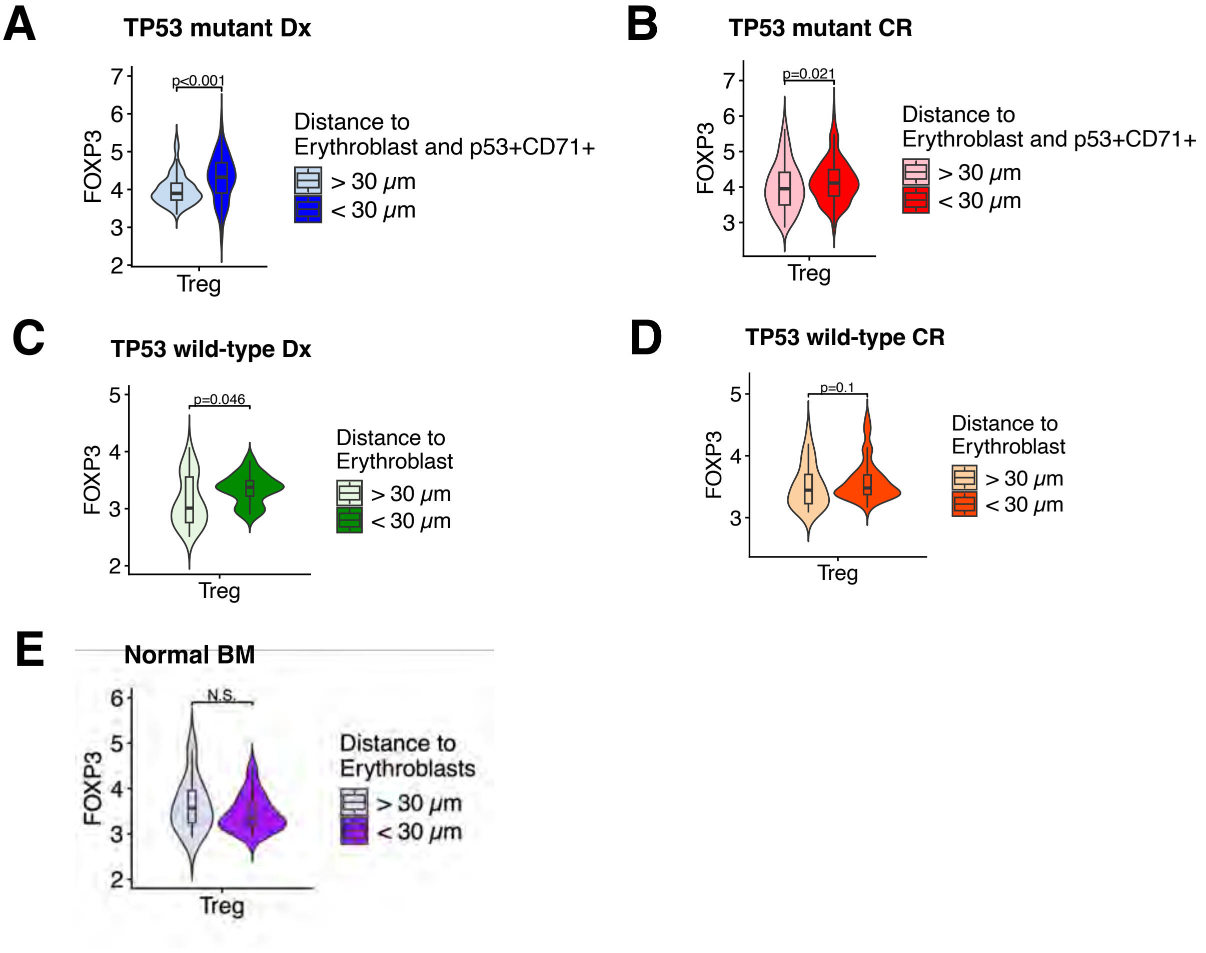

### Supplemental Figure 17

**A**

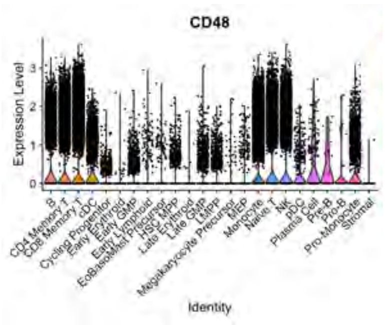


**B**

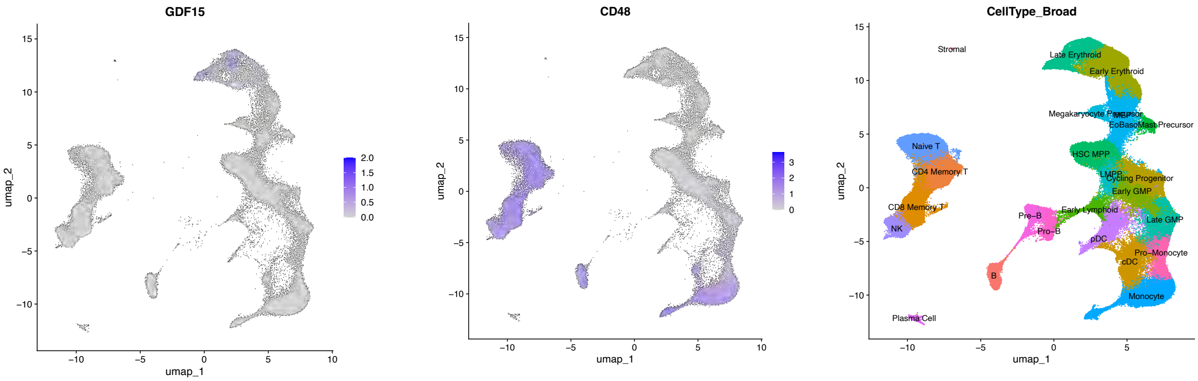


**C**

**Zeng 2025**

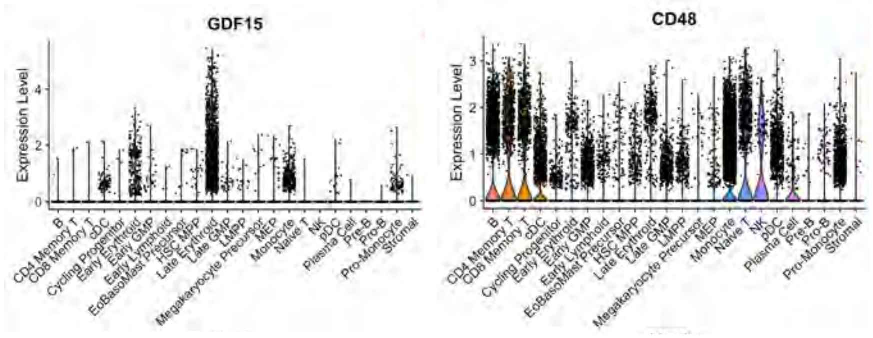


**D**

**van Galen 2019**

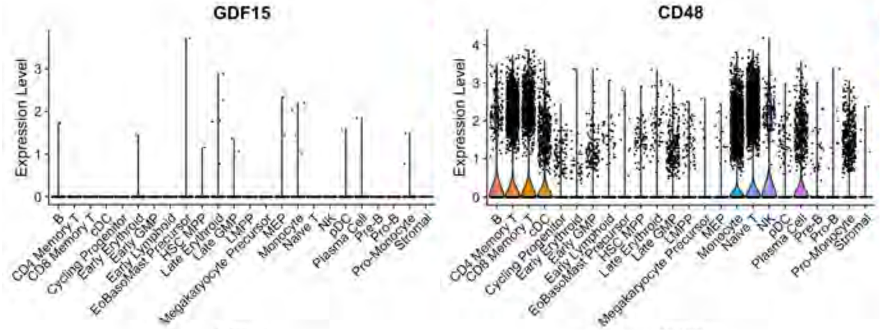

### Supplemental Figure 18

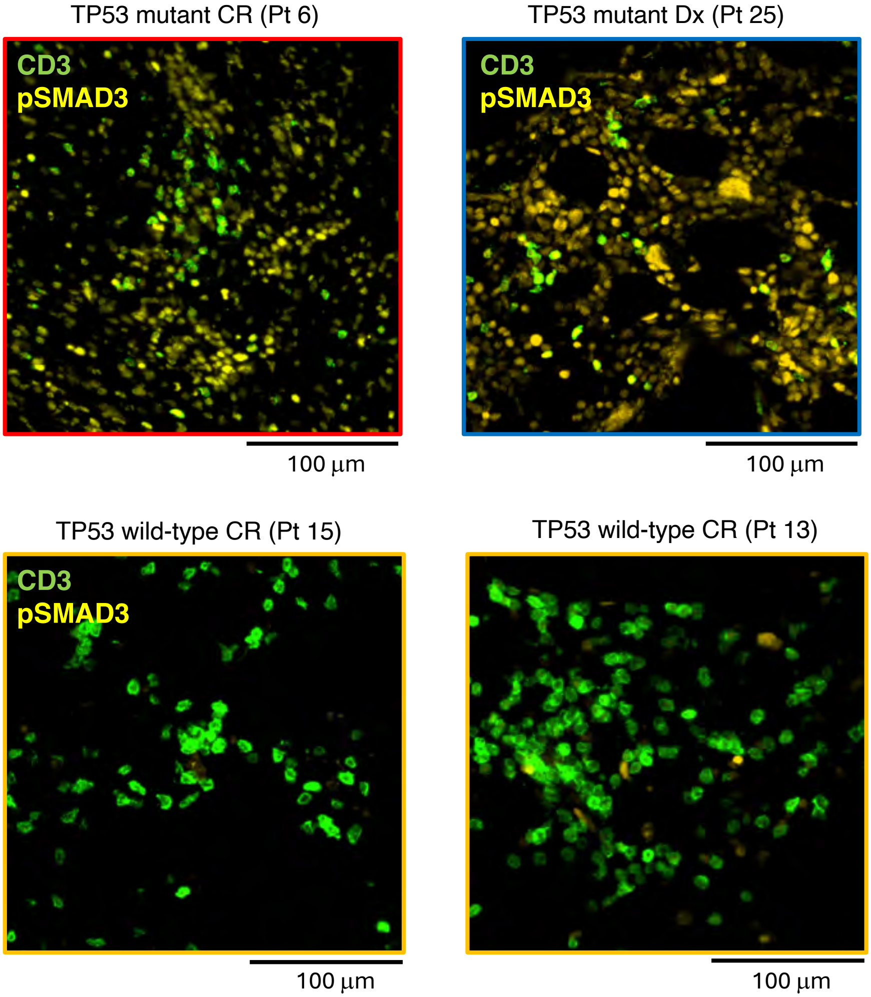

### Supplemental Figure 19

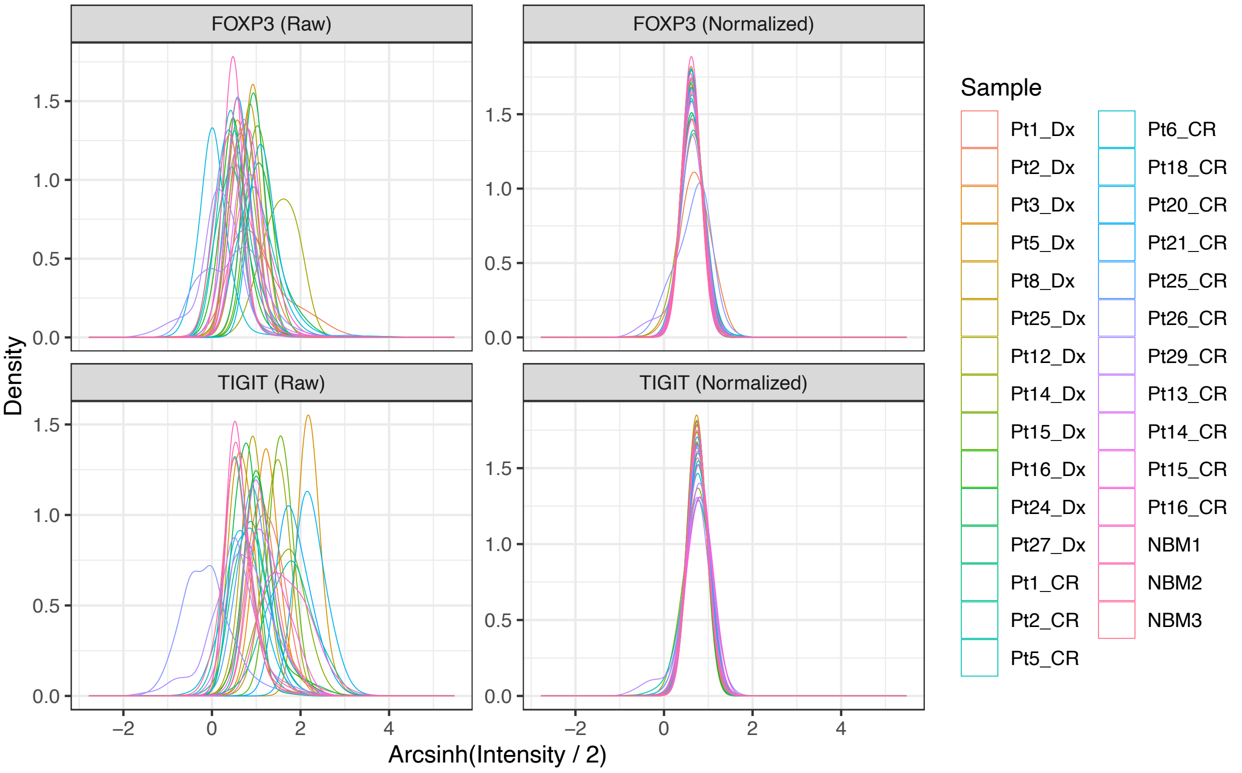
